# Advances on the bacterial diversity and functionality of *Platypus cylindrus - Quercus suber* interaction

**DOI:** 10.64898/2025.12.12.693678

**Authors:** Stefano Nones, Fernanda Simões, Octávio Serra, José Matos, Edmundo Sousa

## Abstract

The ambrosia beetle *Platypus cylindrus* Fab. (Coleoptera: Curculionidae: Platypodinae) has been associated in Portugal with cork oak (*Quercus suber*) tree death since the 1980’s. Although traditionally known as a secondary pest damaging stressed and dead trees, its aggressiveness changed over the last decades. Ambrosia beetle is an insect known for its interactions with several species of fungi aiming for the gain of biologic functions that facilitates its life cycle. How bacterial communities harbored by ambrosia beetle impact forest ecosystems is not yet well known. In oak species, some pathogenic bacteria are often associated with wood insect infestations. From *P. cylindrus* body and wood tree bored galleries, culture dependent methods led to the isolation of bacteria involved in wood degradation and beetle ecology and even a new bacterial cork oak pathogen. To complement colony isolation efforts for the identification of the beetle-cork oak associated bacterial community, the same samples were analyzed with 16S metabarcoding. Two distinct bacterial communities from beetle mycangia and wood galleries sharing the most abundant bacterial taxa were revealed. A group of genera from Enterobacterales and *Bradyrhizobium* prevailed in the samples from wood galleries. Comparison between ASVs and full 16S sequences allowed the identification of Pectobacteriaceae as the major taxon from unclassified Enterobacterales carried by the beetle. Prediction of metabolic pathways varying between wood galleries and beetle mycangia revealed upregulation in the galleries of carbohydrate and virulence factors. Together, these analyses efficiently compared the bacteriome from the wood galleries with the beetle mycangia, revealing different bacterial communities and specific ecological roles related to both the beetle and trees. Additionally, they gave insights into the comparability of culture-dependent and independent methods in bacteriome analysis.

## Introduction

Bacteria inhabit a large set of forest environments ranging from soil to the atmosphere, from living beings to rock surfaces. Forest habitats are distinct, but tightly connected through a network of complex relations (Baldrian, 2017). Bacterial communities contribute to wood decomposition in decaying trees and provide nutrients which can benefit other organisms (Tláskal et al., 2017). Bacteria can also be involved in the promotion of oak diseases, often occurring along with wood insect infestations and causing increasing concern internationally (Poza-Carrión et al., 2008; Brady et al., 2014a, b, 2017; Sitz et al., 2018). This is particularly relevant considering that insect invasions are the major quantitative ecosystem disturbance of forests, with direct consequences on the bacterial communities. Tree leaf loss due to wood boring beetle infestation causes unstable availability of litter which strongly impacts soil microbes and promotes unbalanced plant-microbe interactions (Baldrian, 2017). Larval galleries of the bark-boring beetle *Agrilus biguttatus* are generally present on oaks with acute oak decline (AOD) in British forests (Brady et al., 2017). This decline disease was firstly described in the UK in 2009 (Brady et al., 2017) and found to be promoted by the bacteria *Brenneria goodwinii*, *Gibbsiella quercinecans* and *Rahnella victoriana*. Cases of AOD were also reported in Portugal, Spain, other European countries and Iran (Bene et al., 2025). Within ambrosia and bark beetle communities, it is known that bacteria are involved in insect development and host colonization (Hofstetter et al., 2015; Nones et al., 2021, 2022a). Bacterial taxa potentially feature cellulolytic activity, nitrogen fixation, pathogenesis, fungal growth promotion, nutrients and pheromones production.

The cork oak (*Quercus suber* L.) is an important evergreen species of the Mediterranean Basin from the Fagaceae family. It promotes landscape biodiversity and incorporates atmospheric carbon into its cork layer. Cork is a natural product that supports local economies and the cork industry. Cork oak forests are characterized by low tree density, grassland, pastures, and shrub, and *Q. suber* is the dominant tree species (Tiberi et al., 2016).

In recent decades, oak forests have suffered from a severe process of decline due to biotic and abiotic stresses that impact ecological interactions. The oak pinhole borer *Platypus cylindrus* Fab. (Coleoptera: Curculionidae) is an insect known for its interactions with several species of fungi with different functions in its colonization process, establishment and development (Sousa and Inácio, 2005). It is known as a secondary pest damaging stressed and dead trees, but its aggressiveness changed over the last decades (Tiberi et al., 2016) and its population is promoting tree death within just a few years in Portugal, Spain and Morocco.

*Platypus cylindrus* adults excavate galleries inside tree trunks, which are further colonized by females to inoculate fungal spores. Subsequently, larvae feed and grow on the mycelium. The wood galleries consist of one entry on the bark, which is unique and placed perpendicular to the trunk surface. The first fragment develops for some centimeters into the inside, before undergoing a first ramification from which two branches are excavated through the growth rings of wood. The network of galleries develops further with several secondary ramifications above and below the initial plan, terminating in a dead end where eggs are laid (Sousa and Inácio, 2005).

Recently, the bacterial community resulting from *P. cylindrus* and *Q. suber* interactions in Portuguese cork oak woodlands was investigated with culture dependent (Nones et al., 2020, 2022a; Fernandes et al., 2022) and independent methods (Nones et al., 2021). It was shown that at phylogenetic level, Enterobacterales order dominated cork oak galleries, while Actinobacteria class was predominant in the beetle body and mycangia (Nones et al., 2020, 2022a). By nailing down the study to the differences within the bacterial communities in the mycangia of female and male beetles, it was shown that bacterial taxa possibly associated with tannase activity are linked to males, while some bacteria of Enterobacterales order are linked to females (Nones et al., 2021). Enterobacterales order include known genera related to plant diseases worldwide (Brady et al., 2014b). In the former studies (Fernandes et al., 2022; Nones et al., 2022a), cork oak galleries revealed the presence of *B. goodwinii* and a potential novel species belonging to the Pectobacteriaceae family that caused wilting symptoms on *Q. suber* plantlets. However, in these studies, no association of the bacterial species with the beetle was found by culture-dependent methods.

Here, we carried out 16S metagenomic analysis aiming to complement the efforts done with classical microbiology in *Q. suber* wood galleries and *P. cylindrus* mycangia. Using the same biological samples from the previous culture-dependent study (Nones et al., 2022a), we intended to identify the bacterial taxa and relative abundance from tree gallery-insect interaction, and to disclose metabolic functional prediction, therefore covering any rare or unculturable bacterial species. In doing so, we addressed the following questions:

1. What is the bacterial community resulting from the interaction of the beetle and the cork oak? Are there any similarities between trees and beetles? How specific are the bacterial communities of the wood galleries and the beetle mycangia?
2. Given that galleries are spatially specific for different stages of the beetle (Sousa and Inácio, 2005) and that findings of Ibarra-Juarez et al. (2020) point out a spatio-temporal distribution of bacteria according to the biological life cycle of the beetle, are there significant differences in bacteria at different portions of the gallery?
3. What is the role of the main predicted metabolic pathways for bacteria found in beetle-gallery ecosystem or environment?
4. Lastly, what additional insights can we get by comparing the full 16S sequences from the culture-dependent method of Nones et al. (2022a) with partial amplicon sequence variants (ASVs) from the culture-independent method used in this study?

## Materials and Methods

### Preparation of cork oak gallery samples

A total of five cork oak trees (T1–T5) of similar age (50 years old) and size (diameter at breast height = 120 ± 16 cm), suffering from *Platypus cylindrus* attacks, were selected from a Portuguese cork oak stand in Alentejo (Portugal). The trees were cut down in winter 2018 and were located up to 500 m from each other. After being brought to the lab, they were maintained at a constant temperature and relative humidity (T = 18 °C; R.H. = 70%). Tree gallery samples were obtained by Nones et al. (2022a). Briefly, three galleries were carefully selected and carved out from three trees. Each gallery was made in pieces and grouped into three biological replicates, depending on their depth inside the trunk. The first part comprised only one ramification that was the closest to the entrance (Beginning, I), the second part comprised inner multiple ramifications (Centre, II), while the third part corresponded to the inner end of the gallery (End, III; Figure 1). Only sapwood samples were collected (Table 1). Cork and phloem were discarded since they were considered rich in non-specific mycobiota (Edmundo Sousa’s personal communication). Surface wood from each gallery was scraped into a sterile vial and suspended in extraction buffer (PBS 10 mM, pH 7.2) and incubated overnight at 6 ± 2°C. Afterwards, an aliquot of 500 µl extract from each sample was stored in a vial at – 70 °C. Only the trees T1, T3 and T5 were considered in this metagenomic study, totaling 27 cork oak gallery samples (3 wood samples x 3 galleries x 3 trees; Table 1).

**Figure 1.**
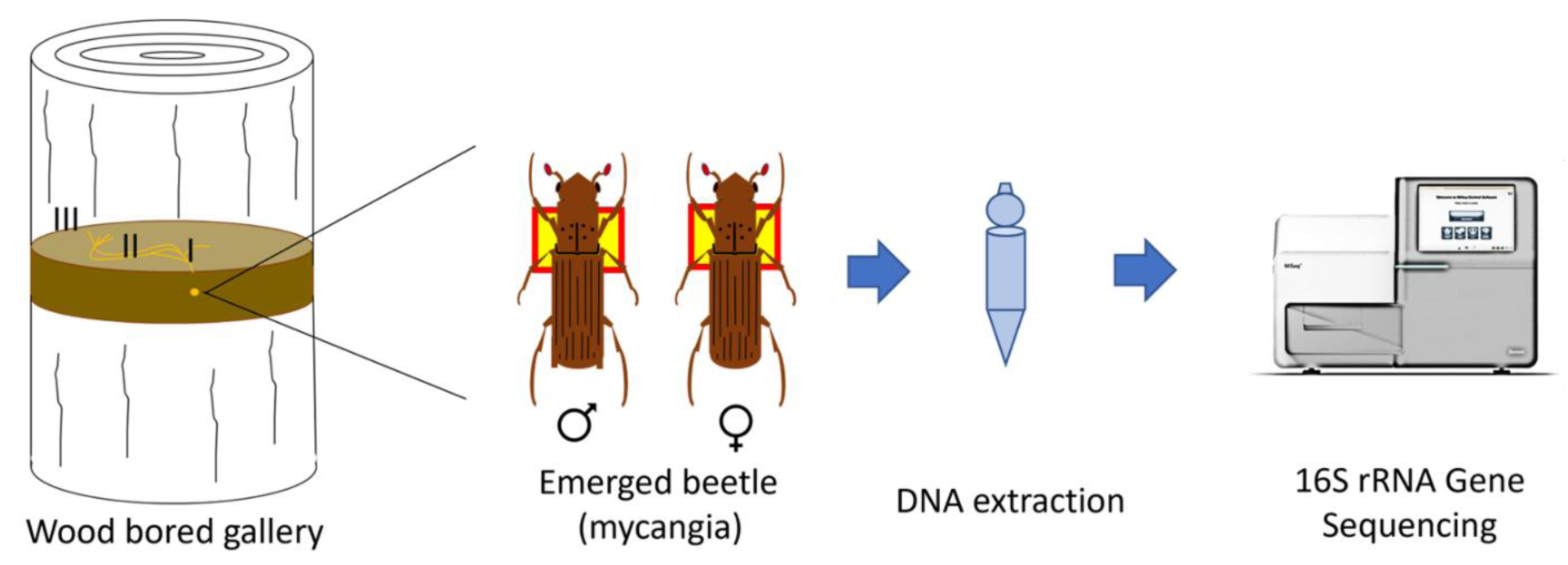
Scheme of the experimental setup of the 16S metagenomic analysis of the wood galleries on *Quercus suber* trees and of the *Platypus cylindrus* beetle mycangia.

**Table 1.** Characteristics of the samples used in this study.

| Tree number | DBH<sup>†</sup> (cm) | Wood gallery parts<sup>‡</sup> | Month of extraction (2018) | Female samples (no.)<sup>¤</sup> | Male samples (no.)<sup>¤</sup> | Mycangia in each sample (no.) |
| --- | --- | --- | --- | --- | --- | --- |
| 1 | 138 | I, II, III | March | 3 | 3 | 3 |
| 2 | 106 | - | - | 3 | 3 | 3 |
| 3 | 137 | I, II, III | April | 3 | 3 | 3 |
| 4 | 110 | - | - | 3 | 3 | 3 |
| 5 | 108 | I, II, III | May | 3 | 3 | 3 |
| Summary |  |  |  |  |  |  |
| 5 | 120±16 | 27 | Mar-May | 15 | 15 | 3 |

<sup>†</sup> DBH = Diameter at breast height
<sup>‡</sup> Beginning (I), Centre (II), End (III)
<sup>¤</sup> Each composite sample is obtained from one gallery out of three distinct galleries

### DNA extraction and sequencing of cork oak gallery samples

DNA extraction from the frozen extract of wood galleries was carried out using innuSPEED Bacteria/Fungi DNA Kit (Analytik Jena, Germany), following the manufacturer’s instructions. Two negative control extractions were made by extracting elution buffer and water independently. Sequencing of the V4 region of the 16S rRNA gene using the barcoded primer set 515F-806R (Caporaso et al., 2011) was carried out at the Genomics Unit of Instituto Gulbenkian de Ciência (Oeiras, Portugal), using a 280-multiplex approach on a 2 × 250 bp PE Illumina MiSeq run. Sequence data have not yet been released. A scheme of the methodology is displayed in Figure 1.

### Bacteriome data from beetle mycangia samples

Beetle samples were obtained from the mycangia of female and male beetles (Figure 1) emerged from the galleries of cork trees (T1-T5), totaling 30 samples (2 sexes x 3 galleries x 5 trees) as shown in Table 1. Bacteriome data from beetle samples, although produced simultaneously with cork oak galleries sequences, were deposited in the European Nucleotide Archive (ENA) at EMBL-EBI under the Accession code PRJEB44517 (https://www.ebi.ac.uk/ena/browser/view/ PRJEB44517) and were previously analyzed (Nones et al., 2021).

### Bioinformatic analysis

The raw paired-end FASTQ reads of the samples of beetle mycangia (PRJEB44517) and wood galleries were retrieved from the ENA database (Accessed on September 1^st^, 2021), and processed using the Quantitative Insights Into Microbial Ecology 2 program (QIIME2, ver. 2021.2, Boylen et al., 2019), with default parameters. DADA2 was used for quality filtering, denoising, paired-end merging, and Amplicon Sequence Variant (ASV) calling, using the plug-in q2-dada2 with the denoise-paired method (Truncation and trimming were set to -p-trunc-len-f 179, -p-trunc-len-r 118, -p-trim-left-f 0, and -p-trim-left-r 0) (Johnson et al., 2008; McKinney, 2010; McDonald et al., 2012; Callahan et al., 2016). Quality filtering was achieved by carrying out reads removal with a Phred quality score (Q) less than 24.5 on average. Moreover, the Naïve Bayes classifier was obtained for the V4 region following the procedure of Robeson (2020) with SILVA 138 SSU NR 99 database and trained in QIIME2 with a q2-feature-classifier fit-classifier-naive-bayes (Pedregosa et al., 2011; Bokulich et al., 2018). Taxonomical classification was determined by matching ASVs against the Silva database (Quast et al., 2012) with a q2-feature-classifier classify-sklearn (Pedregosa et al., 2011). Removal of nonbacterial sequences from the feature table (*i.e.* ASV table) and ASV sequences was performed using filter methods. To visualize the taxonomic composition of the samples, a q2-taxa barplot was used. ASV sequences were aligned using MAFFT (Katoh and Standley, 2013) and used to construct a phylogenetic tree with FastTree 2 (Price et al., 2010). Libraries were then rarefied to the same sequencing depth of (i) 2147 sequences for the cork oak dataset, and (ii) 1260 sequences for the whole analysis (beetle and cork oak). Alpha rarefaction plots were obtained with q2-diversity, using default settings except for “steps” (25) and “max depth” (10000) to test adequacy of sampling. To get specific information from the two datasets, two separate features tables were obtained for the wood galleries and beetle mycangia with the command q2-feature-table filter-samples.

Features having only one entry across the samples and with low abundances (10 for ASVs, 20 for genus) of unrarefied data were filtered out from the tables with ASV sequences and with ASVs collapsed at the genus level (q2-taxa collapse), before ANCOM statistical analysis. Relative abundances (%) were obtained with the command q2-feature-table relative-frequency. The heatmaps with taxa or ASVs were generated by QIIME2 with q2-feature-table heatmap, after log_10_ data normalization. Hierarchical clustering was performed using Euclidean distance and average method. The two datasets (beetle mycangia and wood galleries) were qualitatively compared in a Venn diagram, built with the tool InteractiVenn (www.interactivenn.net; Heberle et al., 2015).

### Functional metabolic prediction of the metagenome

The predicted enzymatic functions of bacterial communities were calculated with q2-picrust2, producing a KEGG Orthology (KO) matrix. Level 3 (L3) KEGG pathways were determined by running the R (R 4.0.3, Core Team) function categorize_by_function (Douglas et al., 2018; 2021) on the KO matrix. The heatmap displaying the metabolic pathways’ abundances in the wood galleries and beetle mycangia was built with the web-based tool MicrobiomeAnalyst (www.microbiomeanalyst.ca; Chong et al., 2020) with the module SDA (Shotgun data profiling), after data filtering with default settings (Low count filter: minimum count 4, prevalence in samples 20%; low variance filter: percentage to remove 10% based on inter-quartile range) and centered log-ratio transformation of the L3-KEGG pathways matrix. The KEGG pathways from ‘Human diseases’ were not displayed in the heatmap, except for ‘Pertussis’ (*i.e. Bordetella pertussis*) and for ‘*Staphylococcus aureus* infection’ that are bacteria related to this environment. Hierarchical clustering was performed using Pearson distance and complete method. Additionally, metagenome functions at sequence level for the ASV matching the taxon Pectobacteriaceae were predicted using the original python implementation of PICRUSt2 v2.5 (https://github.com/picrust/picrust2). Functional categories were mainly built considering relevant biochemical pathways of similar interactions (Broberg et al., 2019; Doonan et al., 2019, 2020; Ibarra-Juarez et al., 2020).

### Statistical analysis

The phylogenetic tree and the rarefied feature table were used to calculate alpha and beta indexes incorporating phylogenetic information. Four alpha diversity indexes (observed features, Shannon, Faith’s Phylogenetic Diversity (PD), Pielou’s evenness) and three beta diversity metrics (Bray–Curtis, Jaccard, and weighted UniFrac) were calculated in QIIME2 using q2-diversity (Lozupone et al., 2005; Lozupone et al., 2007; Chang et al., 2011; Chen et al., 2012; McDonald et al., 2018). Alpha diversity was compared between the different groups of samples using the Kruskal–Wallis test. Principal coordinate analysis (PCoA) was performed based on Bray–Curtis, Jaccard and weighted UniFrac distances and visualized with EMPeror (Vázquez-Baeza et al., 2013). Data separation in the PCoA was tested using the Permutational multivariate analysis of variance (PERMANOVA) or the Permutational analysis of multivariate dispersions (PERMDISP), with *p*-values generated based on 999 permutations. Differences in the abundance of bacteria between galleries’ parts were calculated pairwise using ANCOM test with q2-composition. Furthermore, predicted metagenome functions differentially abundant across wood galleries’ parts and between cork oak galleries and the beetles were inferred by edgeR test with false discovery rate (FDR) correction set at *p* < 0.01 in the web-based tool MicrobiomeAnalyst, module SDA (Chong et al., 2020). A principal component analysis (PCA) based on the L3-KEGG pathways matrix was also generated to display the samples from the wood gallery parts and the beetle mycangia.

### BLAST analysis of full 16S rRNA sequences against ASV

A BLAST analysis was carried out to compare the bacterial sequences obtained when a culture-dependent method was used (Nones et al., 2022a) with those ones of the present study. A total of 71 full-length 16S rRNA gene sequences from culturable strains (MW115318-MW115387, MW346377; Accessed on October 26^th^, 2021) isolated by Nones et al. (2020, 2022a) were retrieved from the NCBI database. These 16S sequences had variable lengths up to 1500 bp, and they are hereinafter referred to as “full 16S sequences.” The bacteria were isolated from the same wood galleries’ and beetle mycangia’ samples used here. To this group, two full 16S rRNA gene sequences were added: one was from a *Curtobacterium* sp. strain from cork oak leaves (Nones et al., 2020) and the other one was from a *Brenneria goodwinii* strain isolated from declining cork oak trees in Portugal (Fernandes et al., 2022). All the full-length 16S rRNA gene sequences were then blasted against the ASV database (V4 region of the 16S rRNA gene) here produced using the software BLAST 2.7.1+ (NCBI, USA). Only sequences matching 100% of identity were recorded.

## Results

To study the bacteriome interactions between wood galleries on *Q. suber* trees and the associated ambrosia beetle *P. cylindrus*, the two datasets (beetle mycangia PRJEB44517 and wood galleries, unpublished) were analyzed together achieving a total of 879,896 reads (645,111 reads from the wood galleries and 234,785 from the beetle mycangia; median: 10,618). The total reads were grouped into 1617 ASVs (297 from the wood galleries and 1,418 from the beetle mycangia) with an average length of 245 bp. The ASVs were shared by 50 samples of up to five cork oak trees: 24 from the wood galleries and 26 from the beetle mycangia, after quality filtering (Supplementary Table S1).

### Baseline survey of cork oak galleries and beetle mycangia bacteriome

The dataset of the cork oak galleries was obtained for the first time, revealing information on the microbial environment surrounding *P. cylindrus* during most of its biological cycle within the tree. Conversely, the baseline survey of the mycangia bacteriome was already presented by Nones et al. (2021). To explore the whole bacterial community within the wood galleries on cork oak, we first sought to identify the differences using alpha- and beta-diversity analysis between each tree. Rarefaction was performed for diversity analyses to a low depth, keeping high levels of biological replication at the cost of sampling depth within single samples (Supplementary Figure S1).

Phylogenetic measures of within sample diversity (alpha diversity) such as the Faith’s PD showed that all trees had a similar bacterial richness. However, the other indices showed that bacterial communities in the cork oak trees T1 and T3 were structurally similar, and significantly different from T5 (Kruskal–Wallis; *p* < 0.05), both qualitatively and quantitatively (Supplementary Table S2). A similar result was mirrored by beta-diversity using abundance weighted UniFrac distance metric and PERMANOVA (Supplementary Table S3).

We further investigated whether the bacterial communities changed within each part of the gallery: beginning (I), centre (II) and end (III). All the samples resulted in similar bacterial richness and evenness. However, the richness was qualitatively (Faith’s PD, observed ASVs) slightly higher in the end than in the other two parts of the gallery (Supplementary Table S4), but not significant (Kruskal–Wallis; *p* < 0.05). Moreover, between samples diversity (Bray-Curtis, Jaccard) highlighted small differences between gallery parts depending on the tree of origin (Supplementary Figure S2; Supplementary Table S5), not significant for PERMDISP (*p* < 0.05). Additionally, the wood gallery parts did not harbor any specific genera or ASVs significantly differentially abundant for ANCOM (*p* < 0.05).

Briefly, a total of 13 bacterial phyla, 17 classes, 47 orders, 86 families, and 137 genera were identified in the wood galleries from the three *Q. suber* trees. Pseudomonadota (formerly Proteobacteria) was the predominant phylum (75 ± 24 %), followed by Actinomycetota (formerly Actinobacteriota,14 ± 17 %), Bacteroidota (formerly Bacteroidetes, 6 ± 13 %), and Bacillota (formerly Firmicutes,4 ± 5 %).

### Bacterial communities in the cork oak galleries and beetle mycangia

*Quercus suber* galleries and *P. cylindrus* beetles mycangia revealed two bacterial communities with distinctive structures, although sharing the most abundant bacterial taxa. The beetle mycangia harbored high bacterial diversity with low abundances, while the wood galleries were dominated by a few taxa with high relative abundance. At higher taxonomic level, bacteria of Enterobacterales order (mean of relative abundance between samples ± SD, 48.21% ± 33.80%) were the most represented within the wood galleries, followed by Rhizobiales (18.65% ± 28.02%) and Micrococcales (8.23% ±1 4.25%), accounting for about the 75% of all the bacterial diversity. Of these three orders, Rhizobiales (13.00% ± 10.76%) was one of those with higher relative abundance also within beetle mycangia. Together with Chitinophagales (11.17% ± 9.11%), these were the only two orders with an individual relative abundance greater than10% in the beetle mycangia (Figure 2A; Supplementary Table S6). At genus level a group of unclassified bacteria of Enterobacterales order (45.86% ± 33.15%) and *Bradyrhizobium* (15.62% ± 28.14%) were the prevailing taxa in the wood galleries. Additionally, within the three major orders of the wood galleries listed above, major taxa included Erwiniaceae (1.02% ± 2.08%), *Shimwellia* (1.14% ± 1.37%), *Ochrobactrum* (1.13% ± 2.34%), *Brevibacterium* (4.04% ± 10.56%) and Microbacteriaceae (1.21% ± 2.98%). Furthermore, only four taxa classified at genus level and present in the whisker plot of the beetle mycangia were absent in the wood galleries (*Hydrotalea*, Comamonadaceae, *Alcaligenes* and *Undibacterium*; Figure 2B-C; Supplementary Table S7).

**Figure 2.**
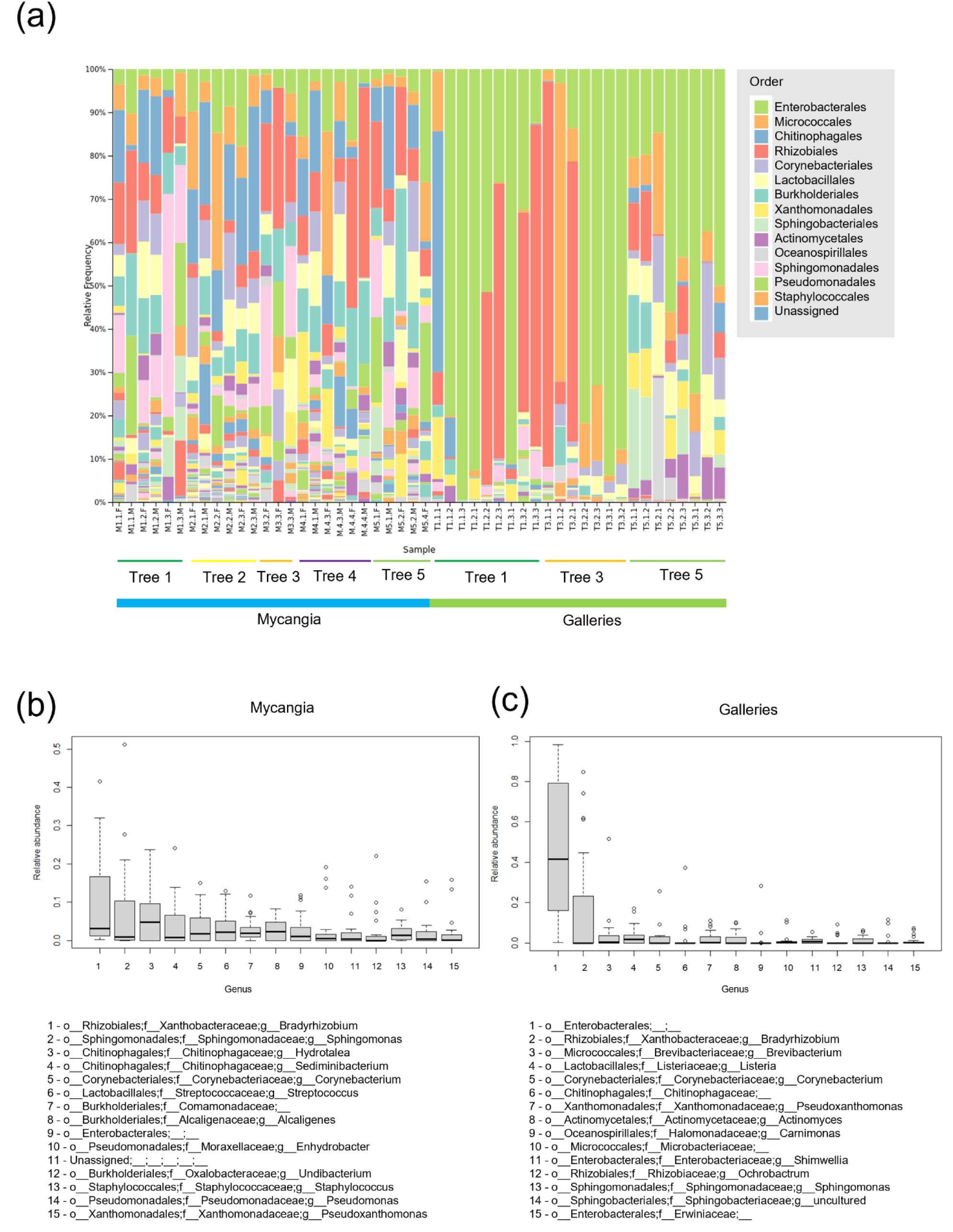
Taxonomic structures of the bacterial microbiota of beetle mycangia and cork oak galleries. **(a)** Taxa bar plot at the order level; **(b**,**c)** Box and whiskers plot of the 15 genera with the largest mean relative abundance across all samples.

From a global perspective, most of the genera involved in this interaction were either only inside the beetle mycangia samples or shared between the wood galleries and the mycangia. In fact, only 24 out of a total of 530 genera were exclusive of the wood galleries (Figure 3; Supplementary Table S8). We therefore investigated the distribution in the wood galleries of the 46 most abundant and frequent genera inhabiting the beetle mycangia, described by Nones et al. (2021) and putatively covering important beetle’s ecological functions. The heatmap indicated that a group of 16 of these bacteria had increased levels also in the wood galleries (Figure 4; Supplementary Table S9). These included unclassified Enterobacterales, *Bradyrhizobium*, *Sphingomonas*, *Corynebacterium*, Comamonadaceae, Microbacteriaceae, Chitinophagaceae, *Ochrobactrum*, *Pseudoclavibacter*, *Actinomyces*, *Nocardioides*, *Pseudoxanthomonas*, *Shimwellia*, Erwiniaceae, *Phyllobacterium* and *Neisseria*. In detail, unclassified Enterobacterales was the only taxon with high relative abundance throughout most of the samples.

**Figure 3.**
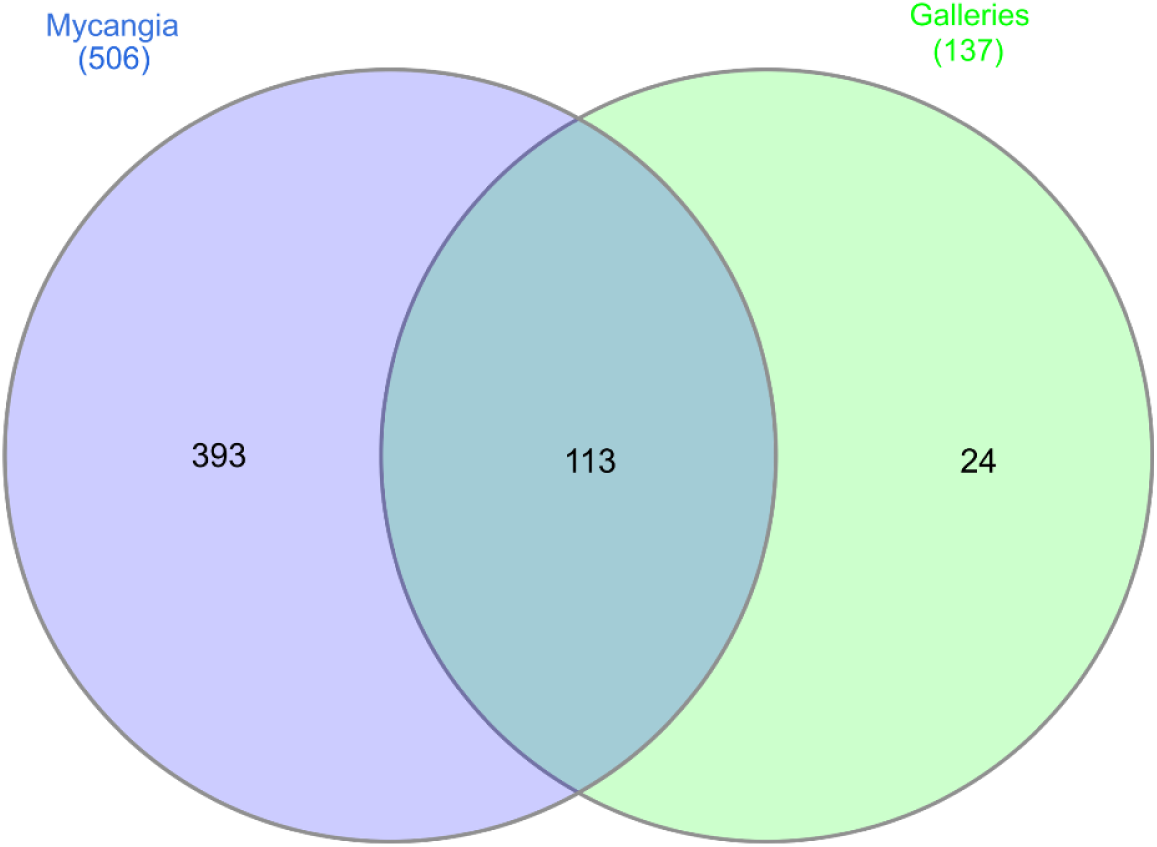
Venn diagram illustrating the genera shared by the beetle mycangia and the wood galleries.

**Figure 4.**
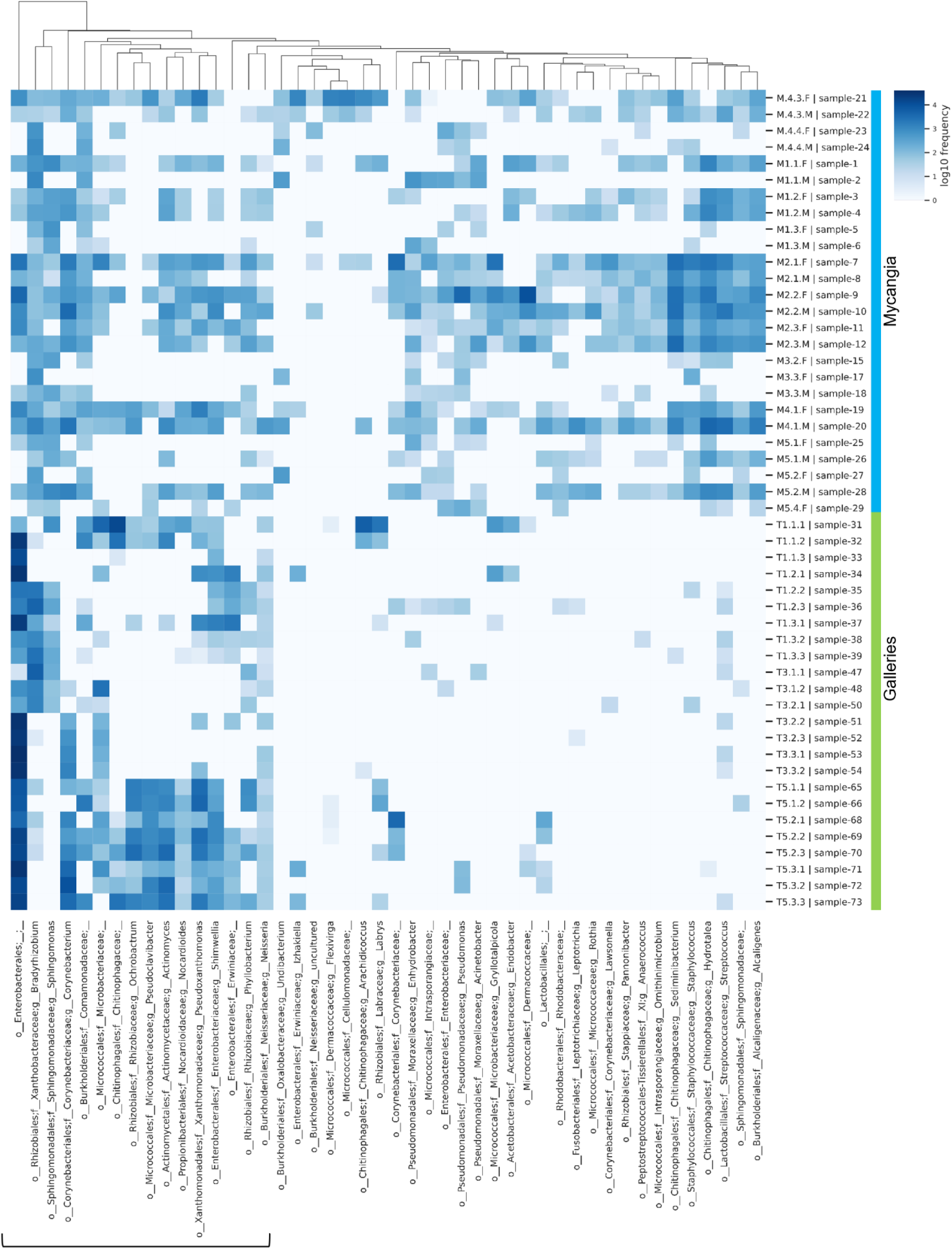
Heat map visualization of the 46 most prevalent and frequent genera (order or family if not further classifiable) inhabiting the beetle mycangia across both mycangia and wood galleries samples, based on normalized mean values. The square bracket marks the genera with increased levels in the wood galleries. Rows: samples; Columns: genera; Color key indicates genera expression value, white: lowest, blue: highest.

### Microbial diversity of cork oak galleries and beetle mycangia

Alpha rarefaction curves (RC) displayed a good coverage of the overall community (Supplementary Figure S1) while maximizing biological replication. RC for Shannon estimator had leveled off, indicating that our analysis covered the quantitative biodiversity within the wood galleries and beetle mycangia. Based on the observed ASV RC, we are aware that a qualitative biodiversity was not revealed for a few samples.

The four indices estimating alpha diversity of the bacteria from the wood galleries reported a much less rich and diverse community than the samples from beetle mycangia (Figure 5a; Supplementary Table S10), statistically significant (Kruskal–Wallis; *p* < 0.05). PCoA based on a weighted UniFrac distance matrix and PERMANOVA showed that beetle mycangia bacteriome was significantly different from the wood gallery ones (Figure 5b; PERMANOVA’s pseudo-F = 11.22, *p* = 0.001). Additionally, the female bacteriome was slightly more similar to the wood galleries than the male bacteriome, although still statistically different from the wood galleries for PERMANOVA (*p* < 0.05; Supplementary Table S11).

**Figure 5.**
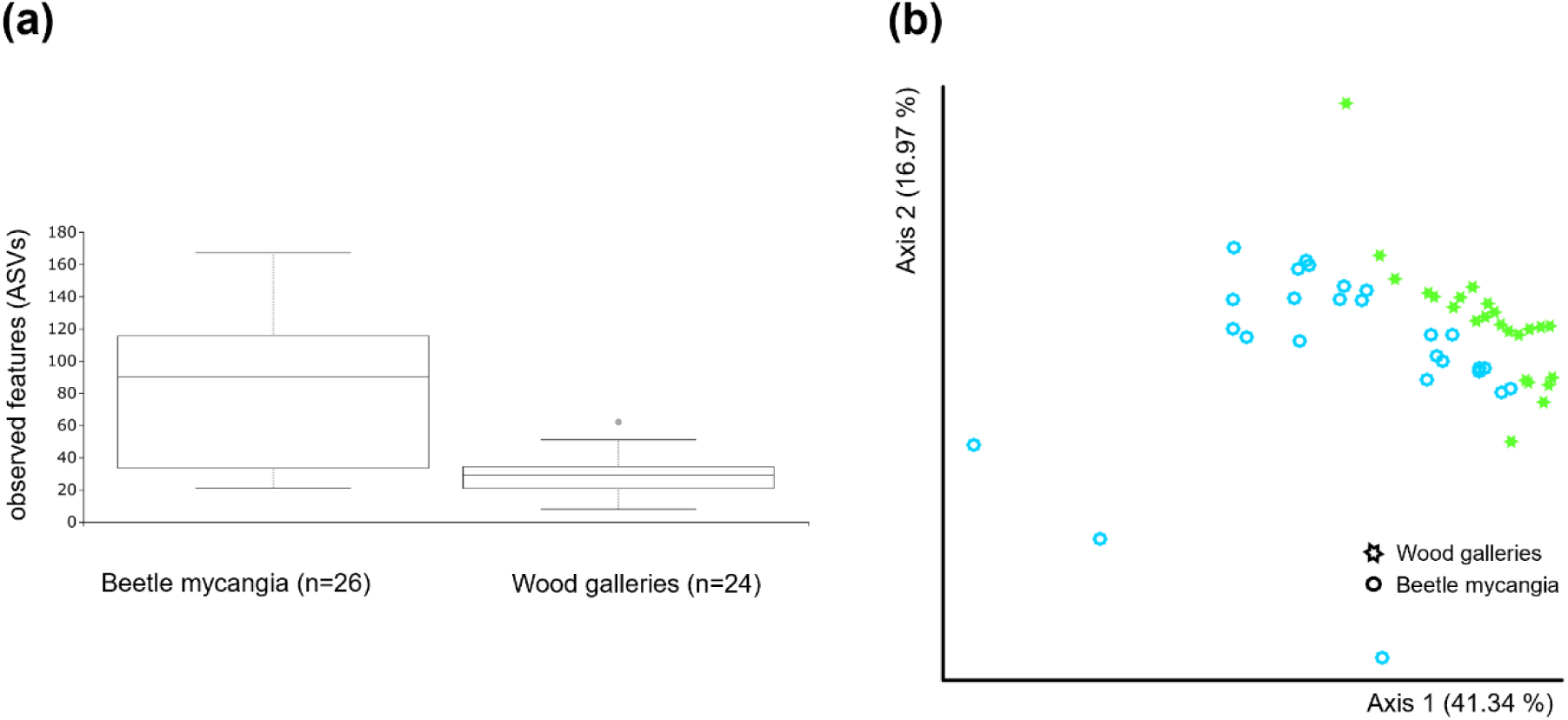
Alpha and beta diversity of beetle mycangia and wood gallery samples. **(a)** Richness (observed ASVs) of beetle mycangia and wood gallery samples represented in box plot; **(b)** Principal coordinate analysis (PCoA) plot based on the weighted UniFrac distance for bacterial communities, showing cluster separation between beetle samples and wood galleries, with significant variation (PERMANOVA’s pseudo-F = 11.22, *p* = 0.001).

### Prediction roles of bacterial communities in wood galleries and beetle mycangia

The predictive analyses of the microbiota in the Kyoto Encyclopedia of Genes and Genomes (KEGG) pathways revealed a total 7061 KEGG orthologs (KOs) grouped into 287 KEGG level 3 (L3) metabolic pathways for bacterial communities of the wood galleries and beetle mycangia. The principal component analysis (PCA) showed that all the samples clustered close to each other in relation to their metabolic pathways (Figure 6). However, samples of the wood galleries were more diverse than the beetle ones, which in contrast clustered the closest, forming an almost private ellipse, as shown in Figure 6. Additionally, four out of six samples from the end of the galleries clustered close to the beetle samples. Most of the samples from the beginning of the galleries were the most distant from the beetle, while samples from the centre displayed an intermediate distance. However, the edgeR test did not show any statistically significant differences among metabolic pathways of bacterial communities across the wood gallery parts, besides for “sporulation” resulting significantly more abundant in the centre of the galleries than at the beginning (edgeR, log2FC=1.41, logCPM =7.72, *p* < 0.05). The heatmap, based on a selection of 46 metabolic pathways differentially abundant (edgeR, FDR adjusted *p*-value < 0.01; Supplementary Table S12) in beetle mycangia and wood galleries, revealed a higher number of significantly more abundant pathways in the wood galleries (Figure 7). Metabolism, especially carbohydrates, was one of these major categories. Moreover, in the wood galleries the pathways related to environmental information and cellular processing were more abundant. Conversely, beetle mycangia showed higher relative abundance for specific pathways involved in antibiotic resistance and biosynthesis (Figure 7; Supplementary Table S13). The top 10 KEGG L3 metabolic pathways with the highest mean relative abundance in the wood galleries are displayed in Supplementary Figure S3.

**Figure 6.**
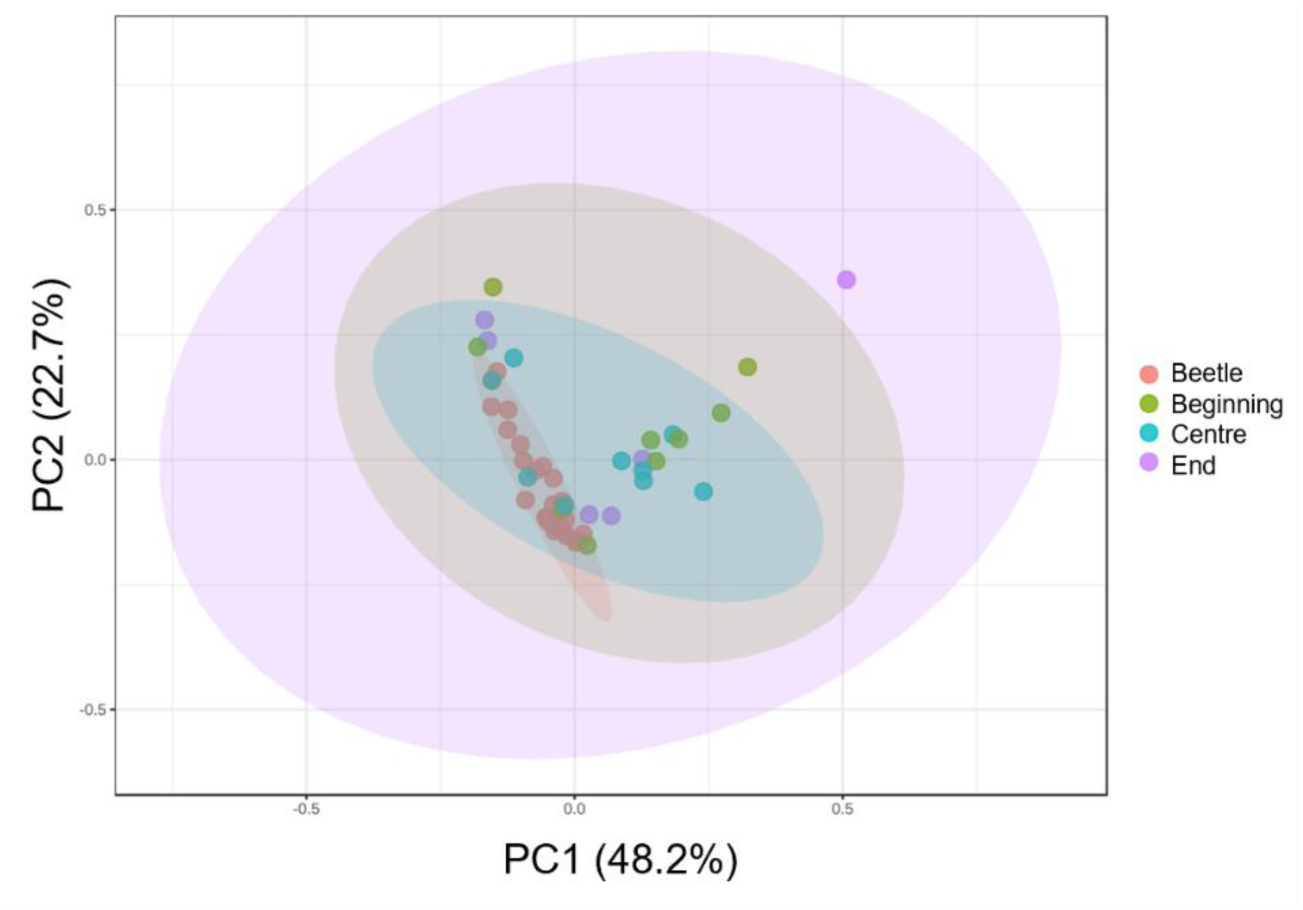
Principal component analysis of predicted KEGG level 3 metabolic pathways of the bacterial community from beetle mycangia and wood galleries. The colored ellipses encircle the samples of the four treatment groups: beetle mycangia, beginning, centre and end of the gallery.

**Figure 7.**
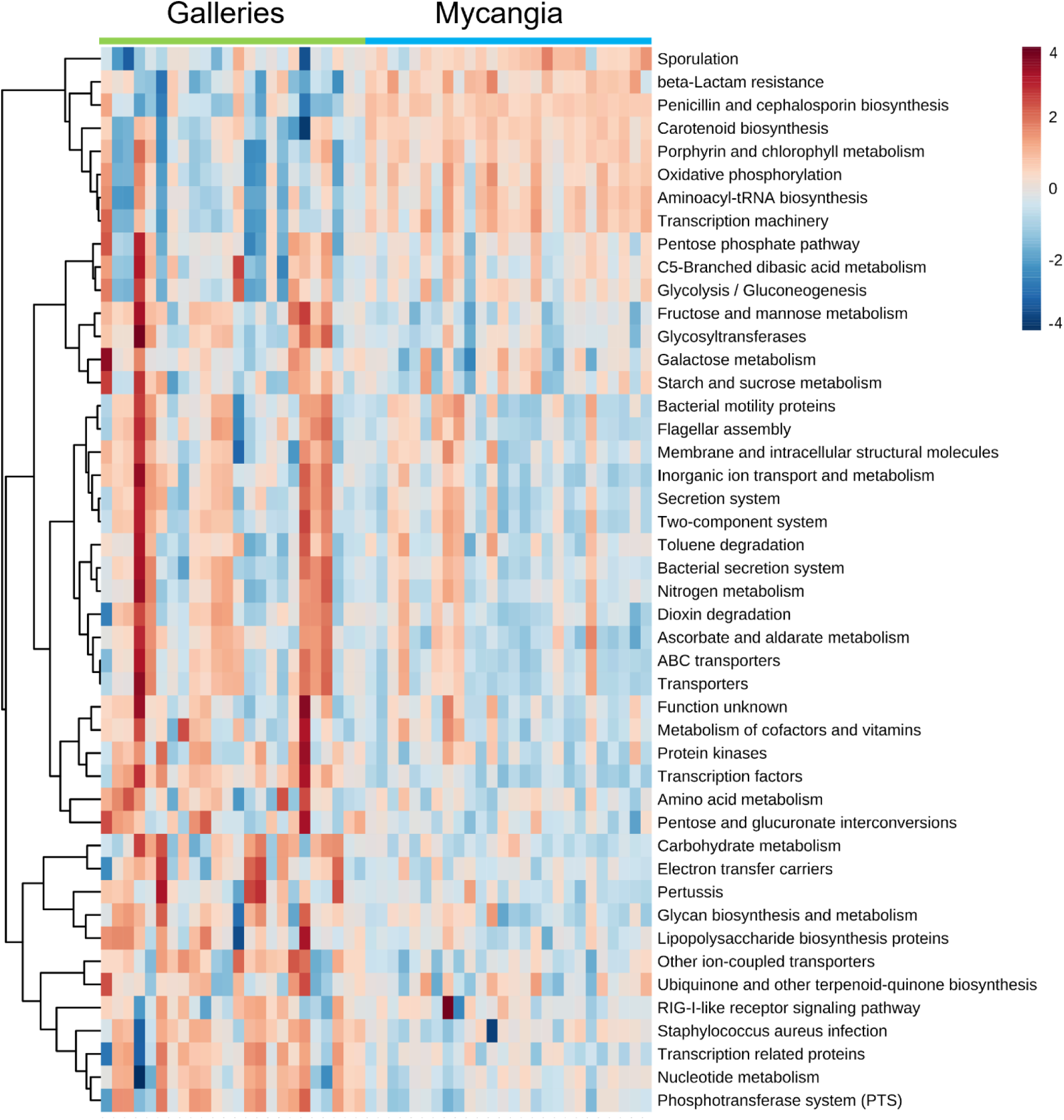
Heat map visualization of 46 predicted KEGG level 3 metabolic pathways differentially abundant across the bacteriome from the beetle mycangia and wood galleries for edgeR statistical test (FDR-adjusted *p* < 0.01). The corresponding KEGG levels 1 and 2 are reported in Supplementary Table S13. Rows: metabolic pathways; Columns: samples; Color key indicates metabolic pathways expression value, white: blue, green: red.

### Complementarity of culture-dependent and independent analyses

Most of the 16S sequences from cultured strains from cork oak galleries and *P. cylindrus* beetles (Nones et al., 2022a), aligned with the sequences from the 16S V4 region from metagenomic analysis when blasted and presenting a match between 99 and 100 % of identity. Moreover, the sequences from the cultured strains identified as *Sphingopyxis* sp. and *Brenneria goodwinii* showed a lower percentage of identity. Conversely, the sequences from one cultured strain of *Niabella* sp., one of Enterobacterales and the cultured strains of *Staphylococcus* spp. had no match at all (Supplementary Table S14).

Taxonomic classification with SILVA database for the 16S sequences from the metagenomic analysis was generally consistent for the matching full-length 16S sequences retrieved from NCBI. Only two sequences had different names. The sequence ASV 10 classified in SILVA as *Pseudoclavibacter* matched *Leucobacter* and ASV 21 classified as *Enhydrobacter* matched *Moraxella*.

The heatmap in Figure 8a displays the variation in relative abundances of the 30 sequences from the V4 region of 16S having 100% of match identity with 52 full-length 16S sequences from cultured strains (Supplementary Table S15). A group of four ASV sequences (ASV 01, 02, 04, 06) classified as Enterobacterales, *Shimwellia* and *Pseudoxanthomonas* showed high variation in both beetle mycangia and wood galleries. Moreover, another group of ten ASVs (03, 05, 07, 09-15) showed high variation mainly in the wood galleries. These were classified in SILVA as Erwiniaceae, Microbacteriaceae, *Brachybacterium*, *Brevibacterium*, Alcaligenaceae, *Gordonia*, *Pseudoclavibacter* and *Ochrobactrum*. Figure 8b shows the matching 16S sequences retrieved from NCBI and their absence/presence adapted from Nones et al. (2022a).

**Figure 8.**
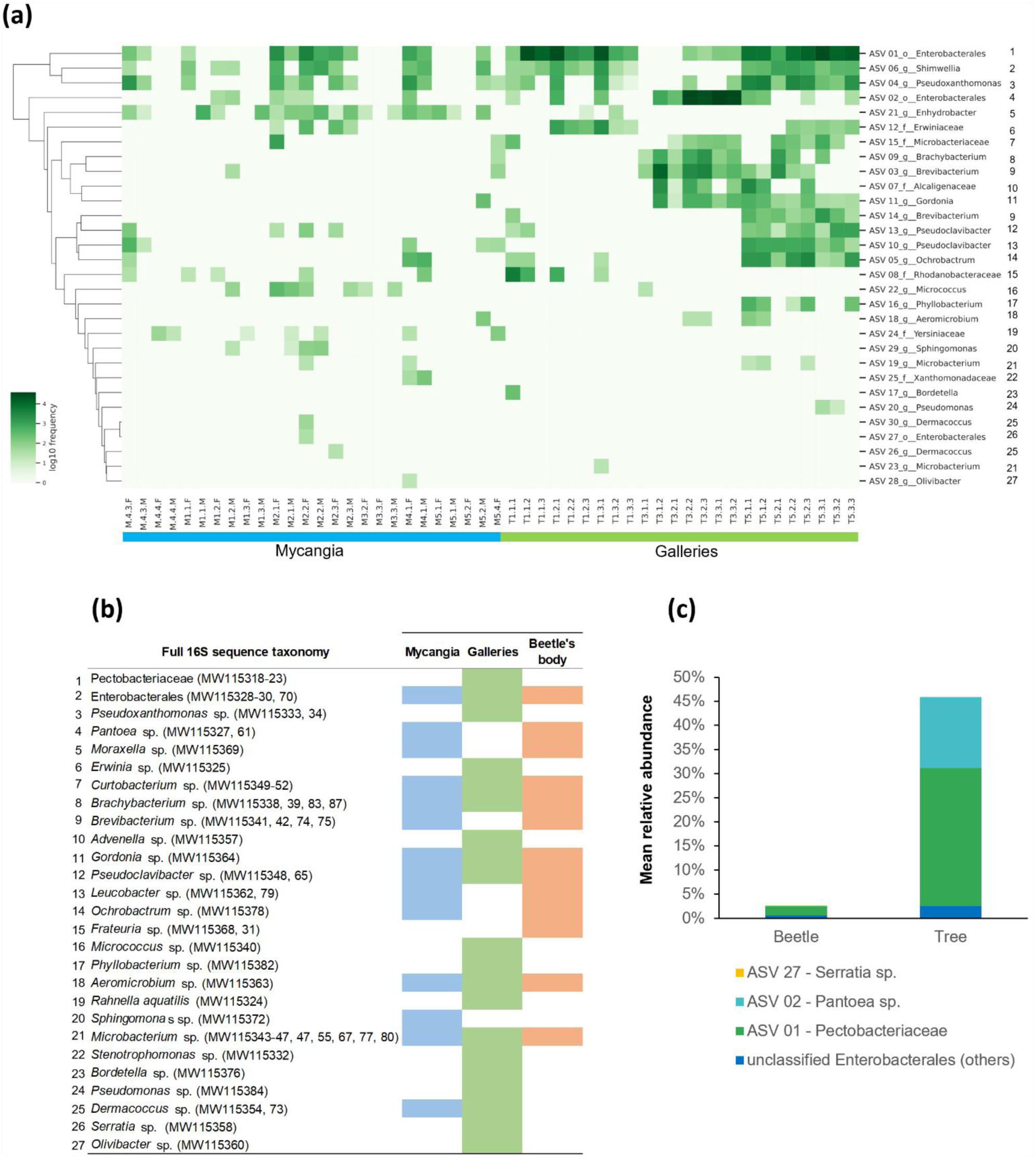
Graphical representation of the matching sequences from full-length and V4 region of 16S rRNA genes obtained from culture-dependent (Nones et al., 2022a) and -independent analyses (this study). **(a)** Heat map visualization of the 30 ASVs classified at genus level with a 100% match with full-length sequences and their relative abundance across beetle mycangia and wood galleries. The column of numbers on the right links the taxonomy of the ASVs with that one of the full-length sequences. Rows: samples; Columns: genera; Color key indicates genera expression value, white: lowest, green: highest; **(b)** Taxonomy of full-length 16S sequences and their presence across beetle and tree samples (adapted from Nones et al., 2022). The column of numbers on the left links the taxonomy of the full-length sequences with that one of the ASVs; **(c)** Stacked Bar plot of the ASV sequences of the taxonomic group unclassified Enterobacterales.

The matches between sequences revealed that the taxonomic group previously classified in SILVA at genus level as Enterobacterales (*i.e.* unclassified) is composed of at least three different taxa: an unknown bacterium of the Pectobacteriaceae family, *Pantoea* sp. and *Serratia* sp. (Figure 8c). Most importantly, the bacterium from the Pectobacteriaceae family accounts for the 28.58% ± 31.04 % (mean of relative abundance between samples ± SD) in the wood galleries and it is the only taxon of this group of Enterobacterales strains present also in the beetle mycangia with a relative abundance over the 0.15% (1.87% ± 3.41 %).

To further investigate the putative metabolic capabilities of the ASV 01 showing high match with the Pectobacteriacae strains, we predicted its KOs and the relative copy numbers. We found that within its 2277 KOs (*i.e.* roughly one third of the KOs in the whole oak-beetle interaction), the ones with more than one copy are mainly related to polar amino acids (*e.g.* aspartic acid, serine, and lysine), methionine, branched-chain and aromatic nonpolar amino acids (amino acids metabolism category, 260 KOs), PCWDEs (plant cell wall degrading enzymes), secretion system, multidrug resistance and defence system (virulence category, 144 KOs), iron and nickel transport (*i.e.* iron ABC transporters and receptors, siderophore receptors like ferrienterochelin and enterobactin, and metalloproteasis like yydH; transition metals category, 82 KOs), and carbohydrates (48 KOs non-PCWDEs and virulence factors; 292 KOs in total (Supplementary Table S16; Supplementary Figures S4-7). Conversely, the lipids and nitrogen cycle (∼40 KOs each) had mostly single copies of KOs.

## Discussion

The present study on the bacterial microbiome associated with the mycangia of *P. cylindrus* beetles and their galleries in the trunk of adult *Q. suber* trees provides a number of novel insights into the bacterial community and respective functional prediction. Primarily, this is the first comparative study addressing the bacteriome resulting from the interaction of a beetle and its host tree showing oak decline symptoms (Sapp et al., 2015, 2016; Denman et al., 2018). Secondly, it addresses without precedent the study of the bacteriome of an ambrosia beetle with its host under semi-field conditions. This adds on to the recent knowledge on microbiome from *Xyleborus affinis* and *Xyleborinus saxesenii* with their galleries obtained artificially (Diehl et al., 2022; Ibarra-Juarez et al., 2020; Cambronero-Heinrichs et al., 2025). Thirdly, it compares results from V4 16S region metabarcoding with sequences obtained after bacterial isolation using a culture dependent method.

### Bacterial communities in the cork oak galleries and beetle mycangia

This study revealed the predominance of Pseudomonadota in galleries and beetles. This taxon is a dominant phylum on wood of both symptomatic and asymptomatic trees from *Quercus* spp., varying in quality of lower taxonomic ranks based on tree age, site and health status (Meaden et al., 2016; Sapp et al., 2016; da Silva 2019). Moreover, Pseudomonadota are an important phylum also in the mycangia of ambrosia beetles (Cambronero-Heinrichs et al., 2025). *Q. suber* galleries and *P. cylindrus* beetles revealed two bacterial communities with distinctive structures. As expected from a previous study (Nones et al., 2021), the mycangia of *P. cylindrus* showed high bacterial diversity with low abundances at order level. This was possibly related to the different environments that the beetles visit. In contrast, wood galleries bacteria were less diverse and a few taxa prevailed, such as Enterobacterales and *Bradyrhizobium* (Rhizobiales), both Pseudomonadota. This qualitative difference in diversity could be partially explained by the inaccessibility of wood galleries. Indeed, the wood galleries are less exposed to the environment than the beetles, and the entry hole is often obstructed by sawdust and excrements or guarded by a male beetle, making it difficult to access (Sousa and Inácio, 2005). Additionally, in the context of trees with acute oak decline (AOD), it is known that the growth of specific bacterial genera within Enterobacterales, such as *Brenneria goodwinii,* gain greater benefit from competitive interaction with the other bacterial species (Brady et al., 2022). Furthermore, gene expression of bacteria such as *B. goodwinii* and *Gibbsiella quercinicans* can be favored in the wood galleries, as a consequence of beetle-microbe-tree interactions (Doonan et al., 2020). In the study of Sapp et al. (2016) on the bacteriome of wood with AOD disease, it was observed that the most diseased samples had elevated levels of a phylotype from Enterobacteriaceae. Moreover, it is known that these bacteria can reside both on oak trees and on several xylophagous beetles, comprising *P. cylindrus* (Bonham; Gathercole et al., 2021).

In this study, most of the bacteria present in the wood galleries on *Q. suber* trees, were shared with the beetle mycangia. Sixteen bacterial genera among the most abundant ones described by Nones et al. (2021) for the mycangia of *P. cylindrus* had increased levels inside the wood galleries. This supports the importance of such bacteria in the context of beetle establishment on the host tree. As recently proposed (Nones et al., 2021), these bacteria might be involved in plant pathogenesis (Enterobacterales, Erwiniaceae), nutrient supplementation (*Bradyrhizobium*, *Sphingomonas*, *Ochrobactrum*), tannins and polysaccharides degradation (*Actinomyces*, *Corynebacterium*, Microbacteriaceae, *Neisseria, Pseudoxanthomonas*), and fungi control (Chitinophagaceae). Additionally, Comamonadaceae, *Nocardioides*, *Pseudoclavibacter* can be found in association with wood insects (Butera et al., 2012; Evtushenko et al., 2015; Ibarra-Juarez et al. 2020) and *Phyllobacterium* with forest trees (Mantelin et al., 2006). Comamonadaceae are bacteria with broad metabolic capabilities (*e.g.* hydrocarbons biodegradation and denitrification; Khan et al., 2002; 2020); *Pseudoclavibacter* are involved in plant polysaccharides degradation (Chase et al., 2016); *Nocardioides* resist extreme conditions, can be mutualistic plant endophytes, and can secrete or degrade secondary metabolites (Evtushenko et al., 2015); and *Phyllobacterium* are generally plant-associated bacteria with antimicrobial and plant growth hormone biosynthesis capabilities (Mergaert and Swings, 2015). Lastly, bacteria from genus *Shimwellia* are not specific to forest environments (*e.g.* cockroaches) and are involved in alcohol (Priest and Barker, 2010; Urano et al., 2015) and vitamin B_12_ production (Brzuszkiewicz et al., 2012).

Although Nones et al. (2021) suggested that the bacteria potentially involved in tannin degradation and linked to the mycangia of male beetles were occupying the cork and phloem of the tree, this hypothesis could not be tested due our previous experimental set up. Future sampling should also take these wood tissues into account.

### Microbial diversity in cork oak galleries and its relationship with the beetle

Most of the diversity indices suggested that the bacterial community was similar in the wood galleries from trees T1 and T3, while increasing in quantity and quality in the smaller tree T5. Interestingly, T5 was a cork oak tree in which the bacterial composition of the beetle mycangia seemed to be sex related (Nones et al., 2021). The increased diversity in the smaller cork oak T5 could be also related to the time of sampling or to other biometric characteristics.

Furthermore, there were no significant differences in alpha and beta diversity, nor in ASV sequences between gallery parts. This might be related to the biological cycle of *P. cylindrus*, which can present mostly larval stages during winter and early spring (Sousa and Inácio, 2005; Belhoucine et al., 2011). This possibly implies specific nutrition requirements that may influence the wood galleries microbiome (Doonan et al., 2020; Ibarra-Juarez et al., 2020). When compared to the beetle mycangia, wood galleries showed a less rich and diverse community, possibly related to the diverse ecological and biological characteristics of the two organisms (Sousa and Inácio, 2005; Meaden et al., 2016; Denman et al., 2018; Doonan et al., 2020; Nones et al., 2021).

### Prediction roles of bacterial communities in cork oak galleries and beetle mycangia

The winter sampling period corresponds to beetles overwintering within the galleries, in which the major developing stage is the larva (Sousa and Inácio, 2005; Belhoucine et al., 2011). Given the high number of larvae in the galleries, it would be expected to have breakdown of complex sugars in the gallery to supply the high energy demand for fungi as nourishment for the progeny (Ibarra-Juarez et al., 2020). A study on oak sapwood tissues infected by *B. goodwinii* showed that many upregulated genes belonged to sugar transport and catabolism (Doonan et al., 2020). Moreover, the data indicated that the Pectobacteriaceae bacterium *B. goodwinii* was stimulated by glucose and xylose rich substrates (Doonan et al., 2020).

In our study we also predicted that carbohydrate metabolism is the category with the highest number of significantly upregulated Level 3 (L3) KEGG pathways in cork oak wood galleries bacteriome when compared to the one of the beetles. Specifically, the wood galleries bacteriome had upregulated levels of 11 pathways, among which glycolysis/gluconeogenesis (*e.g.* for sugars depolymerization), similarly to *B. goodwinii* and *A. biguttatus* eggs when inoculated in live oak logs (Doonan et al., 2020). Also amino acid metabolism, two-component system and lipopolysaccharide biosynthesis proteins were predicted to be upregulated in wood galleries inoculated with *B. goodwinii* when compared with healthy wood logs (Doonan et al., 2020). The lack of healthy wood samples in our study makes it difficult to draw a final conclusion. Interestingly, the wood galleries infested by *P. cylindrus* displayed a bacteriome with significantly upregulated pathways (typically related to virulence-associated genes, Broberg et al., 2018; or to genes potentially affecting plant physiology, Moramarco et al., 2019) when compared to beetle mycangia. These pathways included bacterial secretion system, secretion systems, ABC transporters, transporters, pertussis, bacterial motility proteins and flagellar assembly. Further interest should be given to pathways such as toluene degradation and vitamin metabolism, given the recent knowledge obtained for such *n*-alkyl benzene in *P. cylindrus*-symbiotic ophiostomatoids-infected cork oak seedlings (Nones et al., 2022b) and for the presence of bacteria from the wood galleries putatively involved in nutrients production. Conversely, bacteria from beetle mycangia showed some upregulated pathways involved in antibiotic biosynthesis and resistance, as well as carotenoids biosynthesis. These molecules may potentially be linked to the microbial control of fungal gardens in ambrosia beetles (Aharonowitz et al., 1992; Scott et al., 2008) and to the beetle’s immunology (Tan et al., 2020), respectively. The sequence ASV 01 matching the Pectobacteriaceae taxon highlighted several predicted pathways including virulence factors and relevant functional categories (*e.g.* high copy numbers for carbohydrate metabolism “glycolysis/gluconeogenesis”). Remarkably, this analysis also revealed a variety of pathways involved in the acquisition and modulation of metal ions like iron, which is pivotal for bacterial growth and survival (Chakkour et al., 2024). Although the involvement of such pathways is reported, yet not elucidated, in this pathosystem (*i.e.* iron transporters upregulation in *B. goodwinii* and *G. quercinecans* co-cultures, Doonan et al., 2020; NI-siderophore presence *in* Erwiniaceae, Cambronero-Heinrichs et al., 2023), it is known that other Enterobacterales bacteria such as *Proteus mirabilis* use the metals pools to increase survival rate and modulate the swarming and virulence. This is achieved through sophisticated machinery of siderophores, hemolysins (shared by other plant pathogenic bacteria), metalloproteases, and iron uptake systems (Chakkour et al., 2024). Likewise, the Pectobacteriaceae strains seem to be equipped with similar components, based on the functional prediction analysis. Therefore, this potential feature requires confirmation by future studies of genome sequencing and meta-transcriptomics.

### Complementarity of culture-dependent and independent analyses

The comparison of the amplicon sequence variants (ASVs) from the V4 region of 16S rRNA gene with the complete sequences of culturable bacteria previously identified in this interaction (Nones et al., 2022a), offered a new point of view on the taxonomic assignment, distribution and quantitative importance of these bacteria across all the samples. Interestingly, most of the ASVs of one of the two prevailing taxa of the wood galleries (*i.e.* unclassified Enterobacterales), matched with three groups of complete 16S sequences, identified as Pectobacteriaceae, *Pantoea* sp. and *Serratia* sp.. Among them, Pectobacteriaceae predominated. This is particularly relevant since the bacterial strains from Pectobacteriaceae were isolated only in the wood galleries by Nones et al. (2022a). Our results show that they are also present in the beetle mycangia and, therefore, they might be vectored by *P. cylindrus*. The bacterial strains from Pectobacteriaceae were phylogenetically related to *Brenneria* spp. and *Dickeya* spp., and potentially belong to a new species with tested mild pathogenicity on *Q. suber* plantlets (Nones et al., 2022a). *Pantoea* was phylogenetically related to *P. cedenensis* (Nones et al., 2022a) and it has cellulose degradation potential (Adams et al., 2011). *Serratia* sp. was phylogenetically related to *S. marcescens* (Nones et al., 2022a), a species found both on forest beetles and trees, having insecticidal properties (Qi et al., 2011) and potentially influencing plant metabolism (Vicente et al., 2016).

Four sequences (ASV 01, 02, 04, 06) were represented in the wood galleries and beetle mycangia by high relative abundance and prevalence. These ASVs matched with the complete 16S sequences of Pectobacteriaceae, *Pantoea* sp., Enterobacterales and *Pseudoxanthomonas* sp.. The comparison of partial amplicon sequences with complete 16S sequences, besides generally increasing the taxonomic resolution, showed that caution is still needed. The ASV 06 was classified in SILVA as *Shimwellia*. However, phylogenetic analysis conducted on the matching strains with constitutive genes (*gyrB*, *atpD* and *infB*), showed that the strains could not be allocated to this genus and that their allocation to a specific taxon within Enterobacterales was unclear (Nones et al., 2022a).

Furthermore, the ASV 21 matched with the complete 16S sequences of *Moraxella* sp., which is phylogenetically related to *Enhydrobacter*, the genus classified with SILVA (Kawamura et al., 2012). Both culture-dependent and independent methods confirmed its beetle origin. The bacterium *Enhydrobacter* was reported as an important constituent of the *P. cylindrus* bacteriome (Nones et al., 2021). However, the potential to belong to the genus *Moraxella*, might indicate different ecological roles within the beetle physiology (An, 2007).

A group of 10 ASVs highly abundant in the wood galleries, matched with a group of eleven complete 16S sequences, enabling a more accurate comparison with the bacterial community described with the culture-dependent method (Nones et al., 2022a). This revealed the presence of highly similar ASVs to the complete sequences of *Erwinia* sp. (as Erwiniaceae), *Curtobacterium* sp. (as Microbacteriaceae), *Advenella* sp. (as Alcaligenaceae), and of both *Leucobacter* sp. (as *Pseudoclavibacter*) and *Pseudoclavibacter* sp., in addition to *Brachybacterium* sp., *Brevibacterium* sp., *Gordonia* sp. and *Ochrobactrum* sp.. The presence of *Brevibacterium* sp., *Leucobacter* sp. and *Ochrobactrum* sp. was confirmed also in the wood galleries, and not only in the beetle mycangia as previously shown by the culture-dependent method (Nones et al., 2022a). Most importantly, Nones et al. (2021) showed that some genera from Erwiniaceae were significantly more abundant in the female mycangia of *P. cylindrus* compared to the males. However, a culturable strain of *Erwinia* sp. was isolated only from the wood galleries (Nones et al., 2022a), leaving a potential missing link between the culture-dependent and independent study. This comparison now fills the gap by showing that the ASV 12 detected in both the beetle mycangia and the wood galleries matches with the culturable strain of *Erwinia* sp.. This strain was phylogenetically related to *E. oleae* (Nones et al., 2022a), a bacterium involved in increasing the severity of olive knot disease caused by *Pseudomonas savastanoi* pv. savastanoi (Buonaurio et al., 2015). Moreover, as discussed previously (Chase et al., 2016; Nones et al., 2022a), *Brachybacterium* sp., *Brevibacterium* sp., *Curtobacterium* sp. and *Leucobacter* sp. were linked to carbohydrates degradation. The bacteria from genus *Gordonia* have multiple hosts and they are involved in the production of bioactive compounds and enzymes (Sowani et al., 2018), while *Advenella* sp. is a soil bacterium (Shmareva et al., 2016).

Lastly, a group of 15 ASVs (Positions 15-27 of Figure 8a) showed low relative abundances and prevalence, partially disagreeing with the culture-dependent study. After matching with the complete 16S sequences, the presence of four species was specified: *Frateuria* sp. (as Rhodanobacteraceae), *Rahnella aquatilis* (as Yersiniaceae), *Stenotrophomonas* sp. (as Xanthomonadaceae) and *Serratia* sp. (as Enterobacterales). The inconsistency with the results of the bacterial frequency from the culture-dependent study was probably highly influenced by the use of only three trees, instead of five, as specified in Table 1. An emblematic case is the singleton matching with a culturable strain of *Serratia* sp.. This strain was observed only in the wood galleries of tree T4 (Nones et al., 2022a), which was not included in this metagenomic study. More surprising is the case of *R. aquatilis*, which was reported as being the most frequent culturable bacterial species in the wood galleries (Nones et al., 2022a). In the metagenomic study, the ASV 24 was detected only in the beetle mycangia. Additionally, the highly frequent culturable strains of *Staphylococcus* sp. had no match at all. Several reasons could be implied in the mismatches: either biases of culture-dependent methods, constraints related with 16S V4 region specificity as previously reported (Meisel et al., 2016), or constraints of considering only 100% of identity matches between sequences. The culture-dependent methodology is in fact limited by the number of isolates across multiple samples. Moreover, bacteria have very different growth rate within a community that can be inhibited or favored. Therefore, a certain margin of error could be expected. Conversely, sequences of different lengths might fail to completely align, resulting in matches with low percentages.

None of the ASVs of this study had a high match with the complete sequence of a culturable strain of *B. goodwinii* isolated from a *Q. suber* tree in Portugal (Fernandes et al., 2022). Lower matches were shown for the phylogenetically related strains from Pectobacteriaceae, whose close relations were described in the work of Nones et al. (2022a). This corroborates the results achieved with the cultured-dependent study, in which this plant pathogen was not detected.

Although some of the most important bacterial genera of this interaction were covered by the culture-dependent method, seven relevant genera of the wood galleries and the beetle mycangia failed to be isolated (Nones et al., 2021; 2022a). These comprised the four gram-negative bacteria *Alcaligenes*, *Bradyrhizobium*, *Hydrotalea*, *Sediminibacterium* and the three gram-positive *Actinomyces*, *Corynebacterium* and *Streptococcus*. Major reasons for bacterial ‘culturability’ are related to low prevalence, slow growth (*i.e. Bradyrhizobium*), resistance to cultural media, fastidious growth requirements, presence of bacteriocins, cross-feeding or metabolic cooperation and to bacterial communication requirements (Jordan, 1982; Vartoukian et al., 2010). It is worthy to point out that the optimal growth temperatures and selective media required by these bacteria were generally higher and different from the ones used in the culture-dependent study (Jordan, 1982; Pascual et al., 1995; Van Trappen et al., 2005; Schaal et al., 2006; Qu and Yuan, 2008; Albuquerque et al., 2015; Beck et al., 2020). Additionally, some *Actinomyces* spp. are notoriously difficult to isolate. Indeed, the filamentous and granular structure of these bacterial colonies favors the growth of concomitant bacteria, hindering the production of pure cultures. Moreover, cells are difficult to remove from an agar medium and to lyse effectively (Schaal et al., 2006). In the culture-dependent study, some microbes with mycelial appearance were removed, after meeting such difficulties in the isolation and identification (Stefano Nones’s personal communication).

As expected, the culture-independent method revealed greater numbers and greater richness of bacteria than the culture-dependent one (Nones et al., 2022a). Moreover, the culture-independent method showed that the biggest diversity was harbored by the beetle mycangia and not by the wood galleries, as previously reported. This suggests that the two cultural media used were more prone to cultivate the bacteria from the galleries. However, other factors were probably involved, such as the insufficient quantity of inoculum and specific community ecology interactions between bacterial taxa.

Overall, the use of 16S V4 region ASVs was an important choice for obtaining reliable results from the BLAST alignments with complete 16S sequences. Indeed, ASVs can have differences as little as one nucleotide (Callahan et al., 2017). Moreover, the use of the 138 version of SILVA database (https://www.arb-silva.de/documentation/release-138/; Quast et al., 2012), revealed to be a successful choice for taxonomic classification of ASVs in perspective of the comparison with full 16S sequences classified with NCBI database. SILVA 138 features extensive taxonomic updates, including abundant modifications in the Enterobacterales order. We are confident that future complementation with other databases will improve the classification (Wang and Cole, 2024). Furthermore, the use of ASVs will facilitate the comparison of specific strains across multiple studies, thanks to the unique signatures that characterize these types of sequences (Callahan et al., 2017).

## Conclusions

In this study we carried out 16S metagenomic analysis on *Q. suber* galleries and *P. cylindrus* complementing past efforts done with classical microbiology. Together, these analyses represented an efficient opportunity to link the strength of culture-independent methods to reveal the overall bacterial diversity with those of culture-dependent methods offering high taxonomic power for specific bacteria. We observed a bacterial community that was generally consistent with our previous studies, but we also found new relevant bacterial taxa of this interaction. Additionally, no differences in the community were found between different gallery parts. We unveiled two distinct communities in which the wood galleries had less diversity, most of which were shared with the mycangia. Furthermore, we discovered that *P. cylindrus* transport in their mycangia a group of cork oak phytopathogenic bacteria from Pectobacteriaceae, which predominates in the galleries. Moreover, we found that the cultural media previously used were likely more selective for the bacterial community of the wood galleries, possibly contributing to the distortion of the real bacterial diversity of the tree and the beetle. In order to reduce biased diversity between high-throughput metabarcoding and culturable methods and achieve a better snapshot of the whole bacterial community, we suggest using mixed 16S variable regions and multiple culture media and conditions. Lastly, we provided knowledge supporting the functional role of the bacterial community, showing shared characteristics with the bacteriome from acute oak decline and other ambrosia beetles. These included bacterial pathways related to sugar metabolism from the galleries’ samples and to antibiotics from the beetle mycangia, likely bound to larvae feeding and to the fungiculture of ambrosia beetles. Overall, we provided new knowledge to better understand the establishment of *P. cylindrus* on declining *Q. suber* trees and its escalating threat to cork oak woodlands over the last 40 years. Further studies are planning to perform genome sequencing on Pectobacteriaceae isolates and to identify microbial metabolites of ecological importance in the tree-beetle interaction.

## Acknowledgments

S.N. acknowledges FCT (PD/BD/128400/2017, COVID/BD/151640/2021) and the ITQB NOVA International PhD Program “Plants for Life” (PD/00035/2013) for the PhD grant. Support from the R&D unit GREEN-IT ‘Bioresources for Sustainability’ (Ref. UIDB/04551/2020), and the Instituto Nacional de Investigação Agrária e Veterinária is also acknowledged. The authors thank Leonor Cruz for contribution to the experimental design (INIAV/University of Lisbon) and Conceição Egas for advice on data analysis (Genoinseq/University of Coimbra).

## Supplementary material

Supplementary material associated with this article can be found in the online version on *bioRxiv*.

