## Supplementary Figure S1-S7 for "Advances on the bacterial diversity and functionality of *Platypus cylindrus - Quercus suber* interaction"

(a)

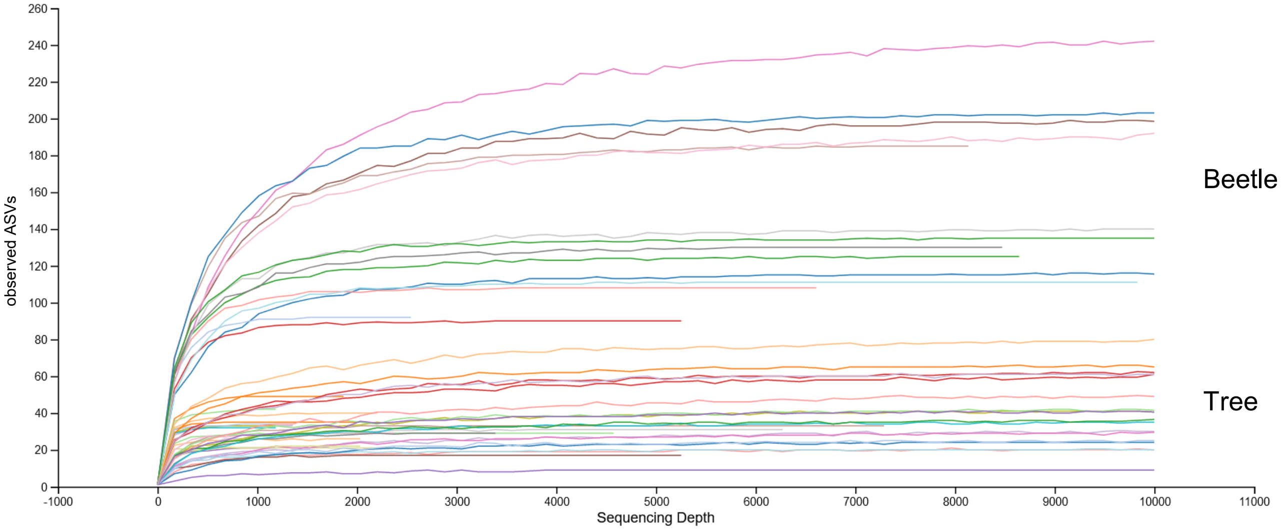

observed ASVs
Sequencing Depth
Beetle
Tree

(b)

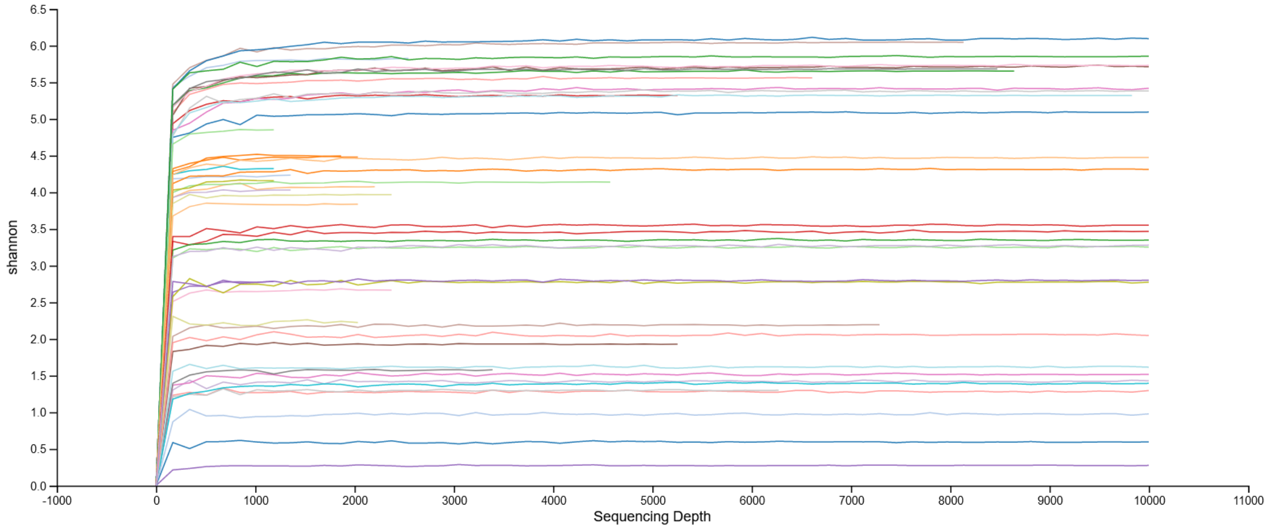

shannon
Sequencing Depth

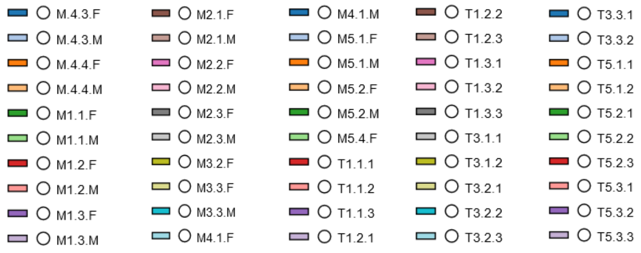

M.4.3.F
M.4.3.M
M.4.4.F
M.4.4.M
M1.1.F
M1.1.M
M1.2.F
M1.2.M
M1.3.F
M1.3.M
M2.1.F
M2.1.M
M2.2.F
M2.2.M
M2.3.F
M2.3.M
M3.2.F
M3.3.F
M3.3.M
M4.1.F
M4.1.M
M5.1.F
M5.1.M
M5.2.F
M5.2.M
M5.4.F
T1.1.1
T1.1.2
T1.1.3
T1.2.1
T1.2.2
T1.2.3
T1.3.1
T1.3.2
T1.3.3
T3.1.1
T3.1.2
T3.2.1
T3.2.2
T3.2.3
T3.3.1
T3.3.2
T5.1.1
T5.1.2
T5.2.1
T5.2.2
T5.2.3
T5.3.1
T5.3.2
T5.3.3

(c)

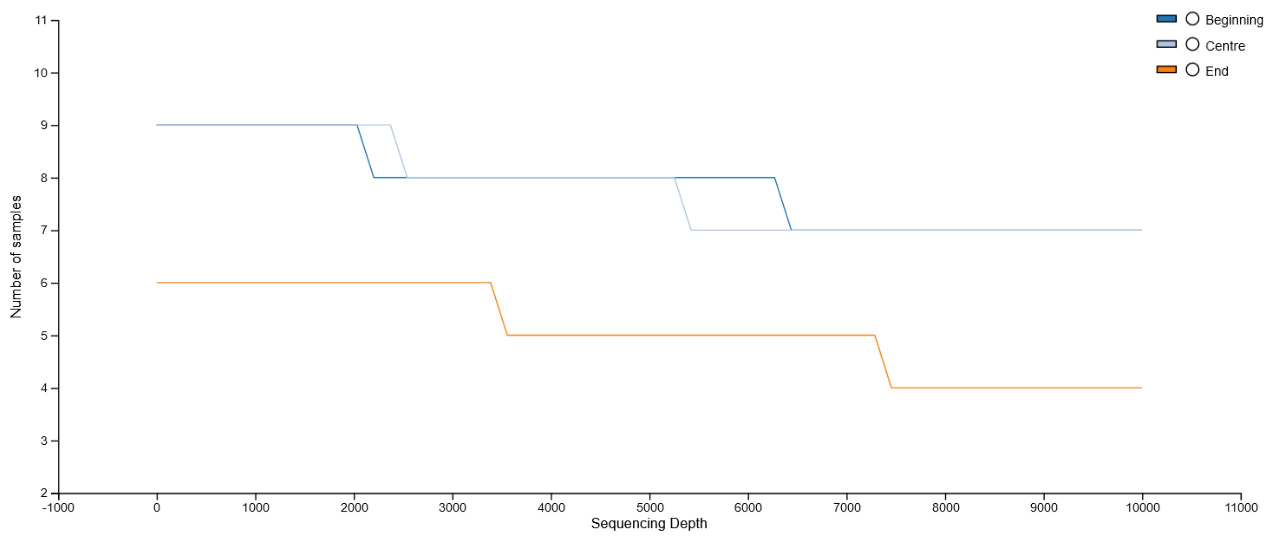

(d)

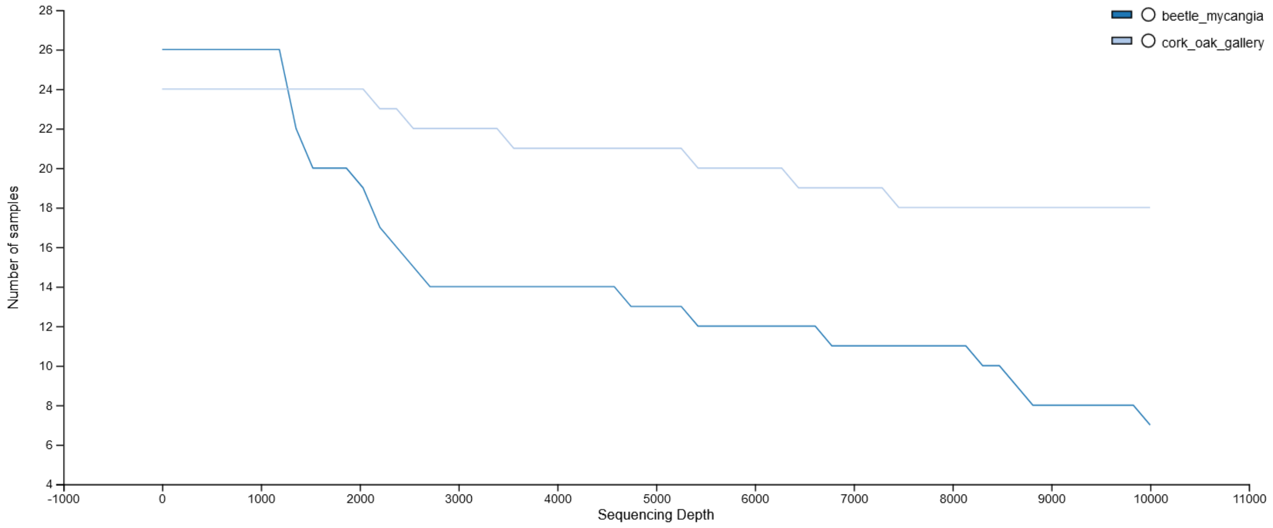

**Figure S1.** Alpha rarefaction curves of the 50 beetle mycangia and wood galleries samples based on high-throughput sequencing (Illumina MiSeq) of bacterial communities: (a) observed ASVs, (b) Shannon, (c) number of samples by gallery's part and (d) number of samples by sample type. Color-coded lines represent all the samples

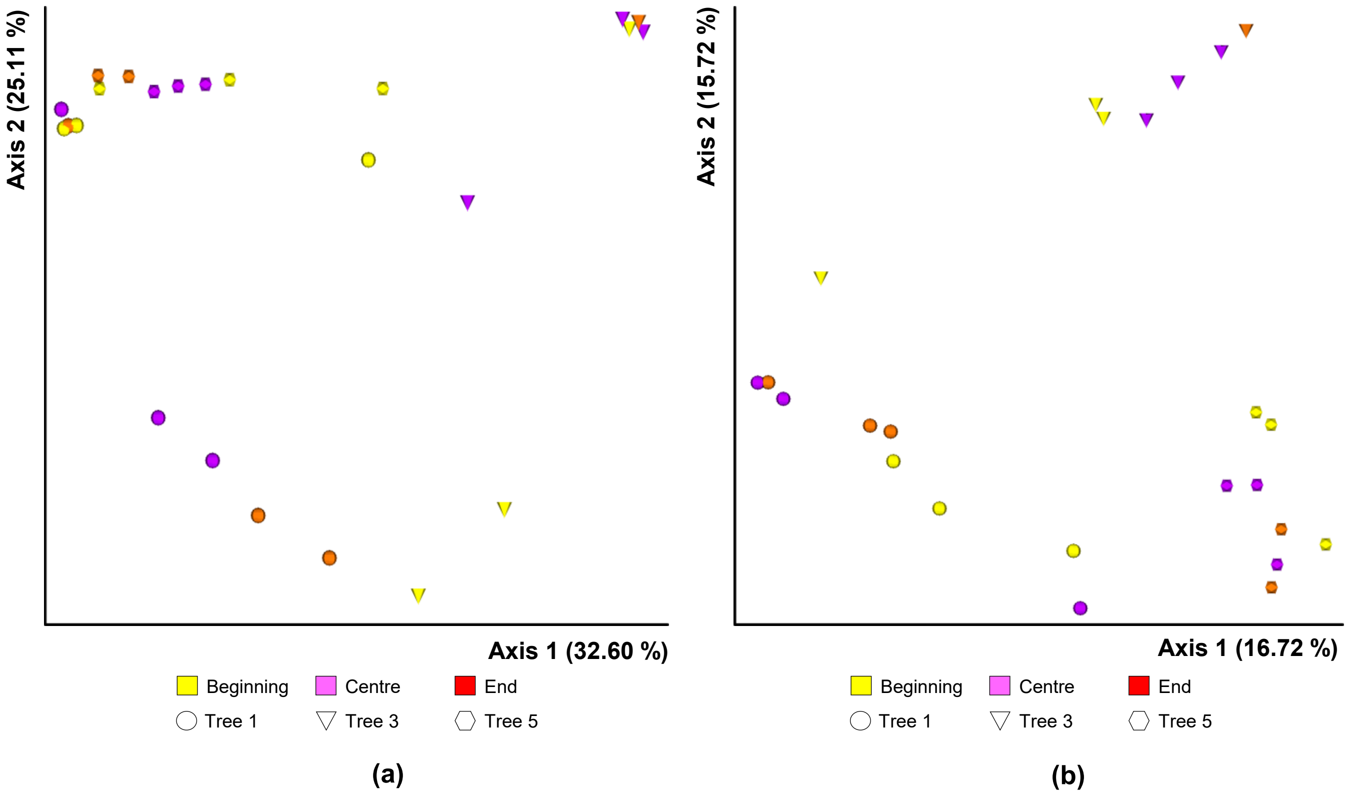

**Figure S2.** Principal coordinate analysis (PCoA) plot based on the (**a**) Bray-Curtis and (**b**) Jaccard distances for bacterial communities, showing cluster separation between wood galleries' parts of individual trees, without significant variation between the beginning, the centre and the end (Bray-Curtis, PERMDISP's pseudo-F = 0.72, *p* = 0.35; Jaccard, PERMIDISP's pseudo-F = 0.11, *p* = 0.88; Supplementary Table S5).

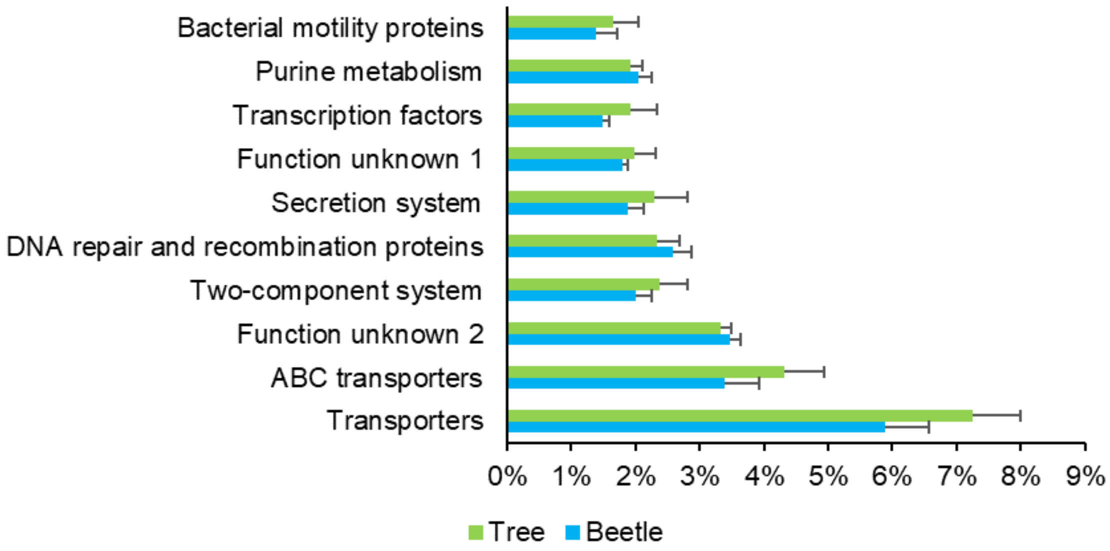

**Figure S3.** Bar charts with the ten majors predicted KEGG level 3 metabolic pathways of the bacteriome from the wood galleries. Mean relative abundances ± SD (%) are displayed for both beetle mycangia and wood-gallery samples.

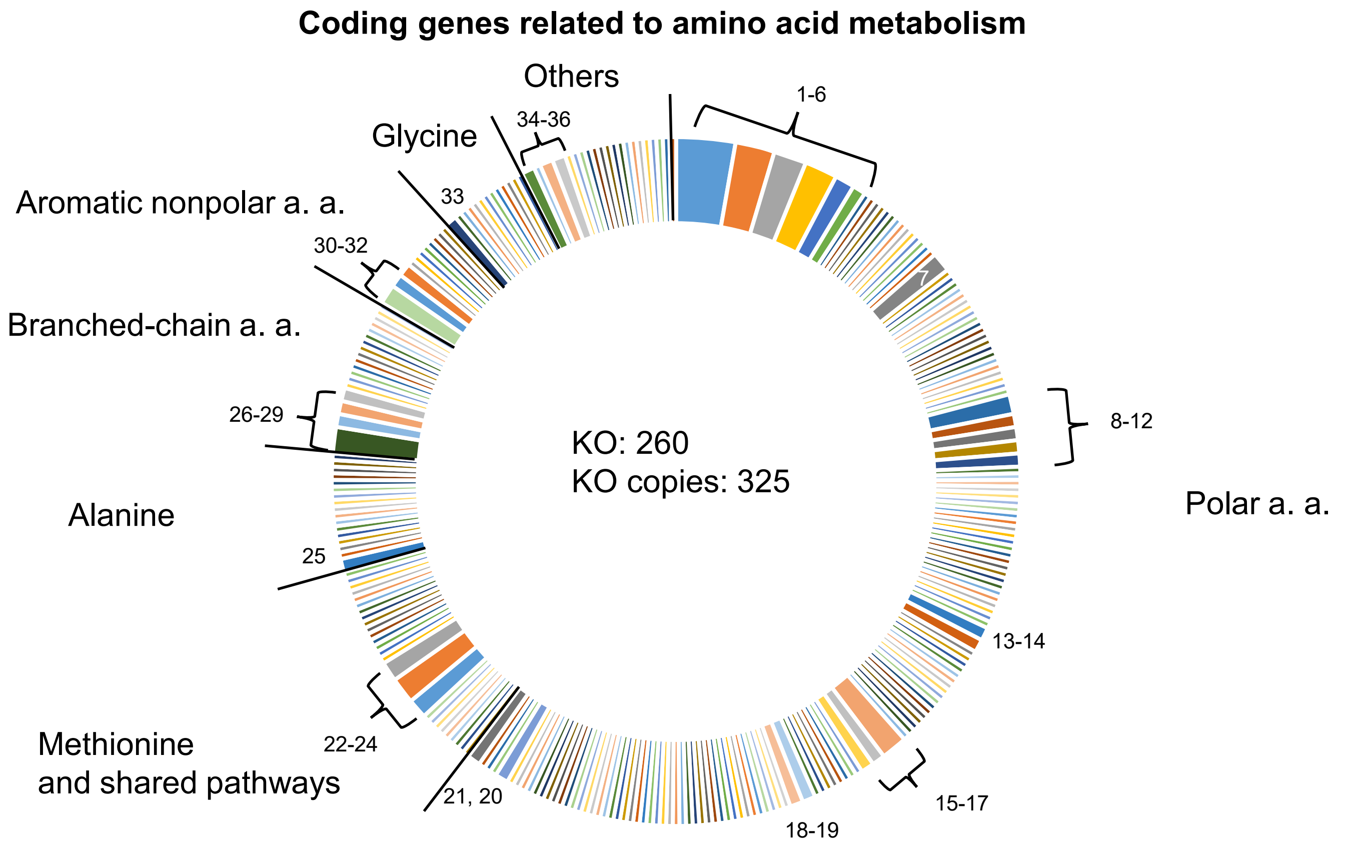

**Legend**

| | |
|---|---|
| 1 | ABC.PA.P; polar amino acid transport system permease protein |
| 2 | ABC.PA.S; polar amino acid transport system substrate-binding protein |
| 3 | ABC.PA.A; polar amino acid transport system ATP-binding protein [EC:7.4.2.1] |
| 4 | tar; methyl-accepting chemotaxis protein II, aspartate sensor receptor |
| 5 | racD; aspartate racemase [EC:5.1.1.13] |
| 6 | aspA; aspartate ammonia-lyase [EC:4.3.1.1] |
| 7 | metE; 5-methyltetrahydropteroyltriglutamate--homocysteine methyltransferase [EC:2.1.1.14] |
| 8 | hipA; serine/threonine-protein kinase HipA [EC:2.7.11.1] |
| 9 | E4.3.1.17, sdaA, sdaB, tdcG; L-serine dehydratase [EC:4.3.1.17] |
| 10 | tsr; methyl-accepting chemotaxis protein I, serine sensor receptor |
| 11 | dacC, dacA, dacD; serine-type D-Ala-D-Ala carboxypeptidase (penicillin-binding protein 5/6) [EC:3.4.16.4] |
| 12 | serA, PHGDH; D-3-phosphoglycerate dehydrogenase / 2-oxoglutarate reductase [EC:1.1.1.95 1.1.1.399] |
| 13 | cysK; cysteine synthase [EC:2.5.1.47] |
| 14 | sufE; cysteine desulfuration protein SufE |
| 15 | dapA; 4-hydroxy-tetrahydrodipicolinate synthase [EC:4.3.3.7] |
| 16 | lysE, argO; L-lysine exporter family protein LysE/ArgO |
| 17 | lysA; diaminopimelate decarboxylase [EC:4.1.1.20] |
| 18 | pdxA; 4-hydroxythreonine-4-phosphate dehydrogenase [EC:1.1.1.262] |
| 19 | E4.3.1.19, ilvA, tdcB; threonine dehydratase [EC:4.3.1.19] |
| 20 | E3.5.1.1, ansA, ansB; L-asparaginase [EC:3.5.1.1] |
| 21 | guaA, GMPS; GMP synthase (glutamine-hydrolysing) [EC:6.3.5.2] |
| 22 | metQ; D-methionine transport system substrate-binding protein |
| 23 | metN; D-methionine transport system ATP-binding protein |
| 24 | metI; D-methionine transport system permease protein |
| 25 | amiABC; N-acetylmuramoyl-L-alanine amidase [EC:3.5.1.28] |
| 26 | 2.2.1.6L; acetolactate synthase I/II/III large subunit [EC:2.2.1.6] |
| 27 | leuA, IMS; 2-isopropylmalate synthase [EC:2.3.3.13] |
| 28 | E2.2.1.6S, ilvH, ilvN; acetolactate synthase I/III small subunit [EC:2.2.1.6] |
| 29 | E2.6.1.42, ilvE; branched-chain amino acid aminotransferase [EC:2.6.1.42] |
| 30 | E2.5.1.54, aroF, aroG, aroH; 3-deoxy-7-phosphoheptulonate synthase [EC:2.5.1.54] |
| 31 | aroP; aromatic amino acid transport protein AroP |
| 32 | hcxB; hydroxycarboxylate dehydrogenase B [EC:1.1.1.237 1.1.1.-] |
| 33 | glyA, SHMT; glycine hydroxymethyltransferase [EC:2.1.2.1] |
| 34 | purB, ADSL; adenylosuccinate lyase [EC:4.3.2.2] |
| 35 | gabD; succinate-semialdehyde dehydrogenase / glutarate-semialdehyde dehydrogenase [EC:1.2.1.16 1.2.1.79 1.2.1.20] |
| 36 | mtnD, mtnZ, ADI1; 1,2-dihydroxy-3-keto-5-methylthiopentene dioxygenase [EC:1.13.11.53 1.13.11.54] |

**Figure S4.** Coding genes inferred from the KEGG database related to amino acid metabolism in the ASV sequence matching the 16S full sequence strains identified as 'Pectobacteriaceae'.
For representation purposes only major pathways with a number of KEGG ortholog (KO) copies over 2 are displayed in the legend. All the other KOs are accessible through the Supplementary Table S16.

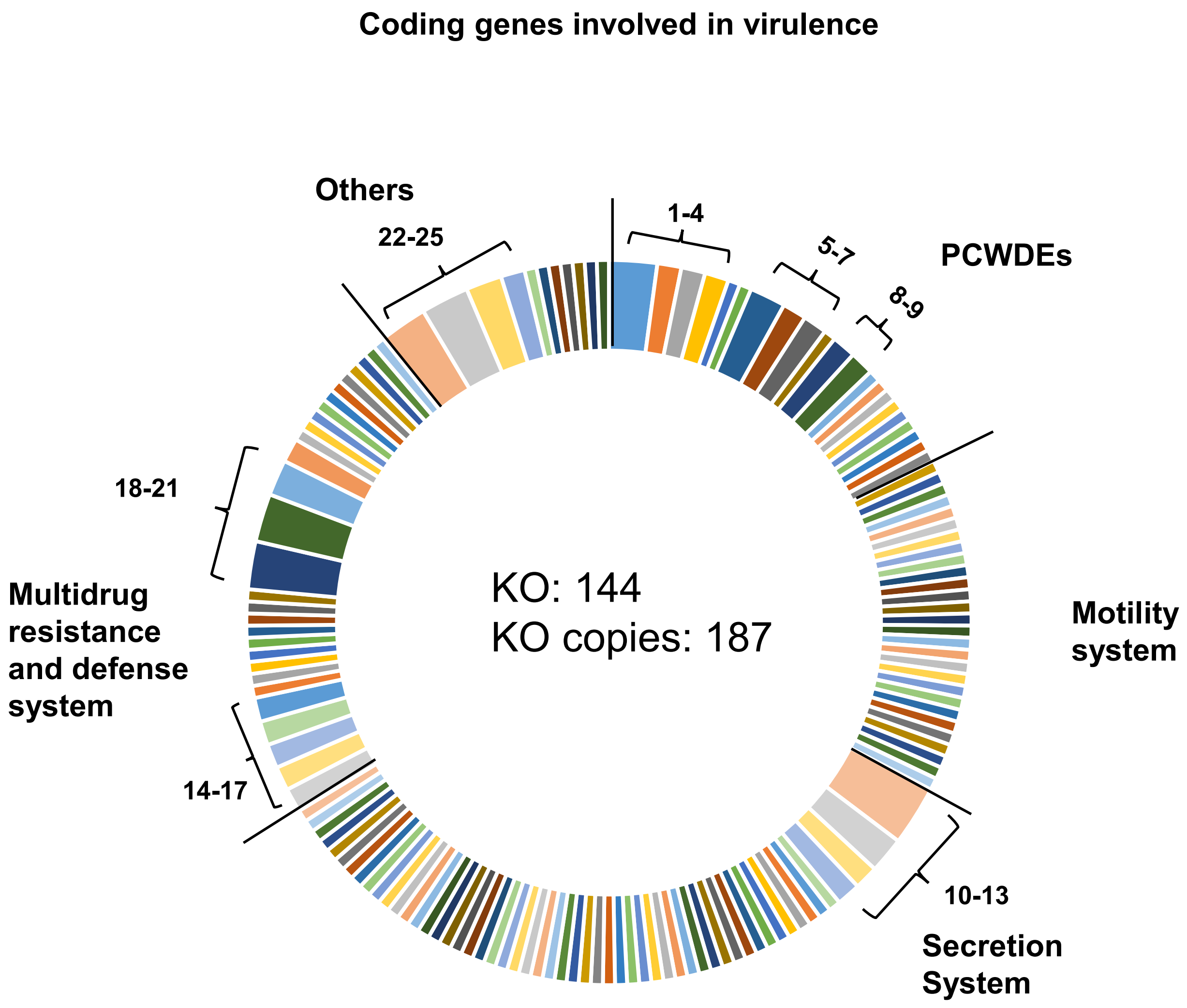

**Figure S5.** Coding genes inferred from the KEGG database related to virulence in the ASV sequence matching the 16S full sequence strains identified as 'Pectobacteriaceae'. For representation purposes only the pathways with a number of KEGG ortholog (KO) copies over 2 are listed: 1 pel; pectate lyase, 2 pectinesterase, 3 uxuB; fructuronate reductase, 4yteR, yesR; unsaturated rhamnogalacturonyl hydrolase, 5 togB; oligogalacturonide transport system substrate-binding protein exuT, 6 MFS transporter, ACS family, aldohexuronate transporter, 7 togT, rhiT; oligogalacturonide transporter, 8 E3.2.1.4; endoglucanase, 9 E3.2.1.89; arabinogalactan endo-1,4-beta-galactosidase, 10 vgrG; type VI secretion system secreted protein, 11 hcp; type VI secretion system secreted protein, 12 ABC-2.TX; HlyD family secretion protein, 13 lapE; outer membrane protein, adhesin transport system, 14 antitoxin CcdA, 15 toxin CcdB, 16 toxin ParE1/3/4, 17 toxic protein SymE, 18 dapA; 4-hydroxy-tetrahydrodipicolinate synthase, 19 oppA; oligopeptide transport system substrate-binding protein, 20 bcr, tcaB; MFS transporter, DHA1 family, multidrug resistance protein, 21 emrE, qac, mmr, smr; small multidrug resistance pump, 22 cspA; cold shock protein, 23 fhaB; filamentous hemagglutinin, 24 trg; methyl-accepting chemotaxis protein III, ribose and galactose sensor receptor, 25 fadA, fadI; acetyl-CoA acyltransferase. All the other KOs are accessible through the Supplementary Table S16.

#### Coding genes related to transition metals

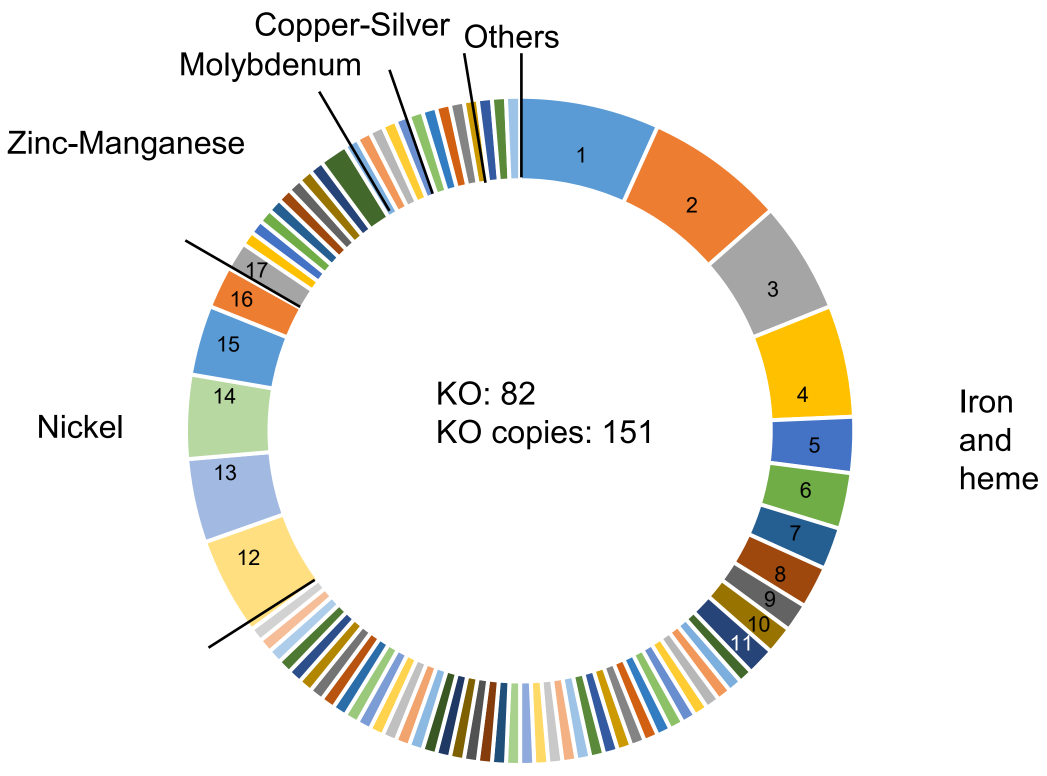

**Legend**

| | |
|---|---|
| 1 | ABC.FEV.P; iron complex transport system permease protein |
| 2 | ABC.FEV.P; iron complex transport system substrate-binding protein |
| 3 | ABC.FEV.A; iron complex transport system ATP-binding protein [EC:7.2.2.-] |
| 4 | TC.FEV.OM; iron complex outermembrane recepter protein |
| 5 | afuC; iron(III) transport system ATP-binding protein [EC:7.2.2.7] |
| 6 | afuB; iron(III) transport system permease protein |
| 7 | afuA, fbpA; iron(III) transport system substrate-binding protein |
| 8 | TC.FEV.OM3, tbpA, hemR, lbpA, hpuB, bhuR, hugA, hmbR; hemoglobin/transferrin/lactoferrin receptor protein |
| | cysG; uroporphyrin-III C-methyltransferase / precorrin-2 dehydrogenase / sirohydrochlorin ferrochelatase [EC:2.1.1.107 |
| 9 | 1.3.1.76 4.99.1.4] |
| 10 | TC.FEV.OM2, cirA, cfrA, hmuR; outer membrane receptor for ferrienterochelin and colicins |
| 11 | fepA, pfeA, iroN, pirA; ferric enterobactin receptor |
| 12 | ABC.PE.S; peptide/nickel transport system substrate-binding protein |
| 13 | ddpF; peptide/nickel transport system ATP-binding protein |
| 14 | ABC.PE.P1; peptide/nickel transport system permease protein |
| 15 | ABC.PE.P; peptide/nickel transport system permease protein |
| 16 | ddpD; peptide/nickel transport system ATP-binding protein |
| 17 | yydH; putative peptide zinc metalloprotease protein |

**Figure S6.** Coding genes inferred from the KEGG database related to transition metals in the ASV sequence matching the 16S full sequence strains identified as 'Pectobacteriaceae'. For representation purposes only the pathways with a number of KEGG ortholog (KO) copies over 2 are displayed in the legend. All the other KOs are accessible through the Supplementary Table S16.

### Coding genes related to carbohydrate metabolism

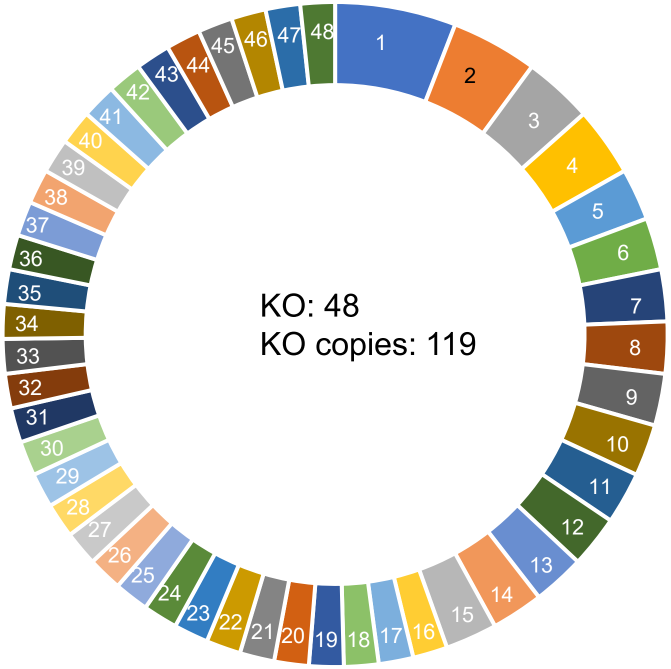

**Legend**

E3.2.1.86B; 6-phospho-beta-glucosidase [EC:3.2.1.86]
galR; LacI family transcriptional regulator, galactose operon repressor celB; cellobiose PTS system EIIC component
E2.2.1.1; transketolase [EC:2.2.1.1]
ABC.SS.P; simple sugar transport system permease protein bglF, bglP; beta-glucoside PTS system EIICBA component [EC:2.7.1.-] celA, chbB; cellobiose PTS system EIIB component [EC:2.7.1.196 2.7.1.205] celC, chbA; cellobiose PTS system EIIA component [EC:2.7.1.196 2.7.1.205] FBA, fbaA; fructose-bisphosphate aldolase, class II [EC:4.1.2.13] frdA; succinate dehydrogenase flavoprotein subunit [EC:1.3.5.1] licT, bglG; beta-glucoside operon transcriptional antiterminator malK, mtlK, thuK; multiple sugar transport system ATP-binding protein [EC:7.5.2.-] rbsC; ribose transport system permease protein
SPP; sucrose-6-phosphatase [EC:3.1.3.24]
ugpB; sn-glycerol 3-phosphate transport system substrate-binding protein ABC.SS.A; simple sugar transport system ATP-binding protein [EC:7.5.2.-] ABC.SS.S; simple sugar transport system substrate-binding protein CS, gltA; citrate synthase [EC:2.3.3.1]
dgoD; galactonate dehydratase [EC:4.2.1.6]
E2.3.1.54, pflD; formate C-acetyltransferase [EC:2.3.1.54] E2.7.1.4, scrK; fructokinase [EC:2.7.1.4]
fruA; fructose PTS system EIIBC or EIIC component [EC:2.7.1.202] gldA; glycerol dehydrogenase [EC:1.1.1.6]
GLO1, gloA; lactoylglutathione lyase [EC:4.4.1.5]
glpA, glpD; glycerol-3-phosphate dehydrogenase [EC:1.1.5.3] glpK, GK; glycerol kinase [EC:2.7.1.30]
gntR; LacI family transcriptional regulator, gluconate utilization system Gnt-I transcriptional repressor hxpA; mannitol-1-/sugar-/sorbitol-6-phosphatase [EC:3.1.3.22 3.1.3.23 3.1.3.50] lacY; MFS transporter, OHS family, lactose permease
pfkA, PFK; 6-phosphofructokinase 1 [EC:2.7.1.11]
pgl; 6-phosphogluconolactonase [EC:3.1.1.31]
PK, pyk; pyruvate kinase [EC:2.7.1.40]
PYG, glgP; glycogen phosphorylase [EC:2.4.1.1]
rbsA; ribose transport system ATP-binding protein [EC:7.5.2.7] rbsB; ribose transport system substrate-binding protein rspA, manD; mannonate dehydratase [EC:4.2.1.8]
TALDO1, talB, talA; transaldolase [EC:2.2.1.2]
TC.GNTP; gluconate:H+ symporter, GntP family
TPI, tpiA; triosephosphate isomerase (TIM) [EC:5.3.1.1] UGP2, galU, galF; UTP--glucose-1-phosphate uridylyltransferase [EC:2.7.7.9] ugpA; sn-glycerol 3-phosphate transport system permease protein ugpC; sn-glycerol 3-phosphate transport system ATP-binding protein [EC:7.6.2.10] ugpE; sn-glycerol 3-phosphate transport system permease protein ulaA, sgaT; ascorbate PTS system EIIC component
ulaB, sgaB; ascorbate PTS system EIIB component [EC:2.7.1.194] ulaC, sgaA; ascorbate PTS system EIIA or EIIAB component [EC:2.7.1.194] xylA; xylose isomerase [EC:5.3.1.5]
xylB, XYLB; xylulokinase [EC:2.7.1.17]

**Figure S7.** Coding genes inferred from the KEGG database related to carbohydrate metabolism in the ASV sequence matching the 16S full sequence strains identified as 'Pectobacteriaceae'. For representation purposes only the pathways with a number of KEGG ortholog (KO) copies over 2 are displayed in the legend. All the other KOs are accessible through the Supplementary Table S16.
